# Diatom pyrenoids are encased in a protein shell that enables efficient CO_2_ fixation

**DOI:** 10.1101/2023.10.25.564039

**Authors:** Ginga Shimakawa, Manon Demulder, Serena Flori, Akihiro Kawamoto, Yoshinori Tsuji, Hermanus Nawaly, Atsuko Tanaka, Rei Tohda, Tadayoshi Ota, Hiroaki Matsui, Natsumi Morishima, Ryosuke Okubo, Wojciech Wietrzynski, Lorenz Lamm, Ricardo D. Righetto, Clarisse Uwizeye, Benoit Gallet, Pierre-Henri Jouneau, Christoph Gerle, Genji Kurisu, Giovanni Finazzi, Benjamin D. Engel, Yusuke Matsuda

## Abstract

Pyrenoids are subcompartments of algal chloroplasts that concentrate Rubisco enzymes and their CO_2_ substrate, thereby increasing the efficiency of carbon fixation. Diatoms perform up to 20% of global CO_2_ fixation, but their pyrenoids remain poorly characterized at a molecular level. Here, we used *in vivo* photo-crosslinking to catalogue components of diatom pyrenoids and identified a pyrenoid shell (PyShell) protein, which we localized to the pyrenoid periphery of both the pennate diatom, *Pheaodactylum tricornutum*, and the centric diatom, *Thalassiosira pseudonana*. *In situ* cryo-electron tomography (cryo-ET) revealed that the pyrenoids of both diatom species are encased in a lattice-like protein sheath. Disruption of PyShell expression in *T. pseudonana* resulted in the absence of this protein sheath, altered pyrenoid morphology, and a high-CO_2_ requiring phenotype, with impaired growth and reduced carbon fixation efficiency under standard atmospheric conditions. Pyrenoids in mutant cells were fragmented and lacked the thylakoid membranes that normally traverse the Rubisco matrix, demonstrating how the PyShell plays a guiding role in establishing pyrenoid architecture. Recombinant PyShell proteins self-assembled into helical tubes, enabling us to determine a 3.0 Å-resolution PyShell structure. We then fit this *in vitro* structure into an *in situ* subtomogram average of the pyrenoid’s protein sheath, yielding a putative atomic model of the PyShell within diatom cells. The structure and function of the diatom PyShell provides a new molecular view of how CO_2_ is assimilated in the ocean, a crucial biome that is on the front lines of climate change.

## Introduction

Diatoms are one of the most dominant groups of phytoplankton in the ocean. They are responsible for 15-20% of the annual global primary production (Falkowski et al., 1998; Smetacek, 1999), powering the Earth’s carbon cycle and feeding energy into vast marine food webs. Despite their importance, the underlying molecular mechanisms that enable diatoms to efficiently assimilate CO_2_ remain poorly understood. Diverse clades of eukaryotic algae, including diatoms, rely on a biophysical CO_2_-concentrating mechanism (CCM) to thrive in CO_2_-limited aquatic environments. Algal CCMs use HCO_3_^-^ transporters to actively accumulate dissolved inorganic carbon (DIC) in the chloroplast and then use carbonic anhydrases (CAs) to convert this HCO_3_^-^ into a high local concentration of CO_2_ in the pyrenoid, a chloroplast subcompartment packed with the carbon-fixing enzyme ribulose 1,5-bisphosphate carboxylase/oxygenase (Rubisco). The pyrenoid thereby floods Rubisco with its CO_2_ substrate, while suppressing Rubisco’s competitive oxygenase reaction, enabling rates of carbon fixation that exceed those of land plants (Giordano et al., 2005; Tsuji et al., 2017; Hennacy and Jonikas, 2020; Shimakawa et al., 2023).

Pyrenoids are a general feature of algal CCMs. However, these chloroplast subcompartments have convergently evolved numerous times and exhibit a wide variety of morphologies (Meyer et al., 2017; Barrett et al., 2021; Uwizeye et al., 2021), indicating that pyrenoids in different clades may have distinct components and specialized mechanisms. To date, only the pyrenoid of the freshwater green alga *Chlamydomonas reinhardtii* has been characterized in molecular detail. The *C. reinhardtii* pyrenoid consists of a spherical matrix of densely packed Rubisco complexes, surrounded by a starch sheath and fenestrated by a network of membrane tubules (Griffiths, 1970; Lacoste-Royal and Gibbs, 1987; Engel et al., 2015). These tubules contain an α-type CA that produces a source of CO_2_ at the center of the pyrenoid (Funke et al., 1997; Raven, 1997; Karlsson et al., 1998; Hanson et al., 2003). The matrix is formed by the liquid-liquid phase separation of Rubisco with its linker protein EPYC1 (Mackinder et al., 2016; Freeman Rosenzweig et al., 2017; Wunder et al., 2018; He et al., 2020), and it dynamically condenses or disperses in response to changes in CO_2_ concentration (Ramazanov et al., 1994; Morita et al., 1997; Borkhsenious et al., 1998).

In contrast to the *C. reinhardtii* pyrenoid, the Rubisco matrix of the diatom pyrenoid has an elongated oval shape and is typically traversed along its long axis by one or two specialized thylakoids (Jenks and Gibbs, 2000; Bedoshvili et al., 2009; Flori et al., 2017). Several proteins have been localized to the pyrenoid of the marine diatom, *Phaeodactylum tricornutum*. In addition to Rubisco, this pyrenoid contains β-type CAs (Tachibana et al., 2011), fructose 1,6-bisphosphate aldolases (FBAs) (Allen et al., 2011), and a θ-type CA specifically localized in the lumen of the thylakoids at the center of the pyrenoid (Kikutani et al., 2016; Shimakawa et al., 2023). Although these observations strongly suggest that diatom pyrenoids increase CO_2_ concentration around Rubisco in a similar fashion to green algae, the pyrenoid components in *P. tricornutum* have distinct origins and arose from endosymbiotic red algae, stramenopile host cells, or diatom-specific bacterial gene transfer (Allen et al., 2011; Nonoyama et al., 2019; Kroth and Matsuda, 2022). In the other words, the pyrenoids of diatoms and green algae may have convergently evolved similar functions from a different set of proteins.

In this study, we identify and characterize a distinct component of diatom pyrenoids that is not present in *C. reinhardtii*: the pyrenoid shell (PyShell). This proteinaceous sheath tightly encases the Rubisco matrix, is required for establishing pyrenoid architecture, and is essential for efficient CO_2_ assimilation and cell growth. We directly observe PyShells in both pennate diatoms (*P. tricornutum*) and centric diatoms (*Thalassiosira pseudonana*), while bioinformatic analysis suggests that PyShells are common in several clades of marine algae, and thus, likely play a major role in driving the ocean’s carbon cycle.

## Results

### Identification and localization of diatom PyShell proteins

To identify novel components of diatom pyrenoids, we performed *in vivo* photo-crosslinking (Suchanek et al., 2005), then disrupted the cells and looked for proteins that co-migrated with the Rubisco large subunit (RbcL) by sucrose density gradient centrifugation and SDS-PAGE (diagrammed in Figure 1A). *P. tricornutum* cells were fed with ʟ-photo-leucine and ʟ-photo-methionine, synthetic amino acid derivatives with diazirine rings in their side chains. Because they are structurally similar to natural amino acids, these photo-reactive amino acids (pAAs) are taken up by the cells and incorporated during protein synthesis. These *P. tricornutum* cells were then irradiated with UV light, causing the pAAs to form reactive carbenes that enable zero-distance photo-crosslinking with directly interacting proteins. Crude extracts from the photo-crosslinked cells were separated by SDS-PAGE and immunoblotted for RbcL (“Procedure A”, Figure 1B). In only the sample with both pAAs and UV irradiation, we observed a RbcL-positive band trapped in the stacking gel, which we analyzed by liquid chromatography tandem mass spectrometry (LC-MS/MS). In an alternative approach, the crude extracts were separated by sucrose density gradient followed by SDS-PAGE (“Procedure B”, Figure 1C). The sample treated with both pAAs and UV irradiation showed RbcL bands in denser sucrose fractions, which we subjected to LC-MS/MS analysis.

**Figure 1.**
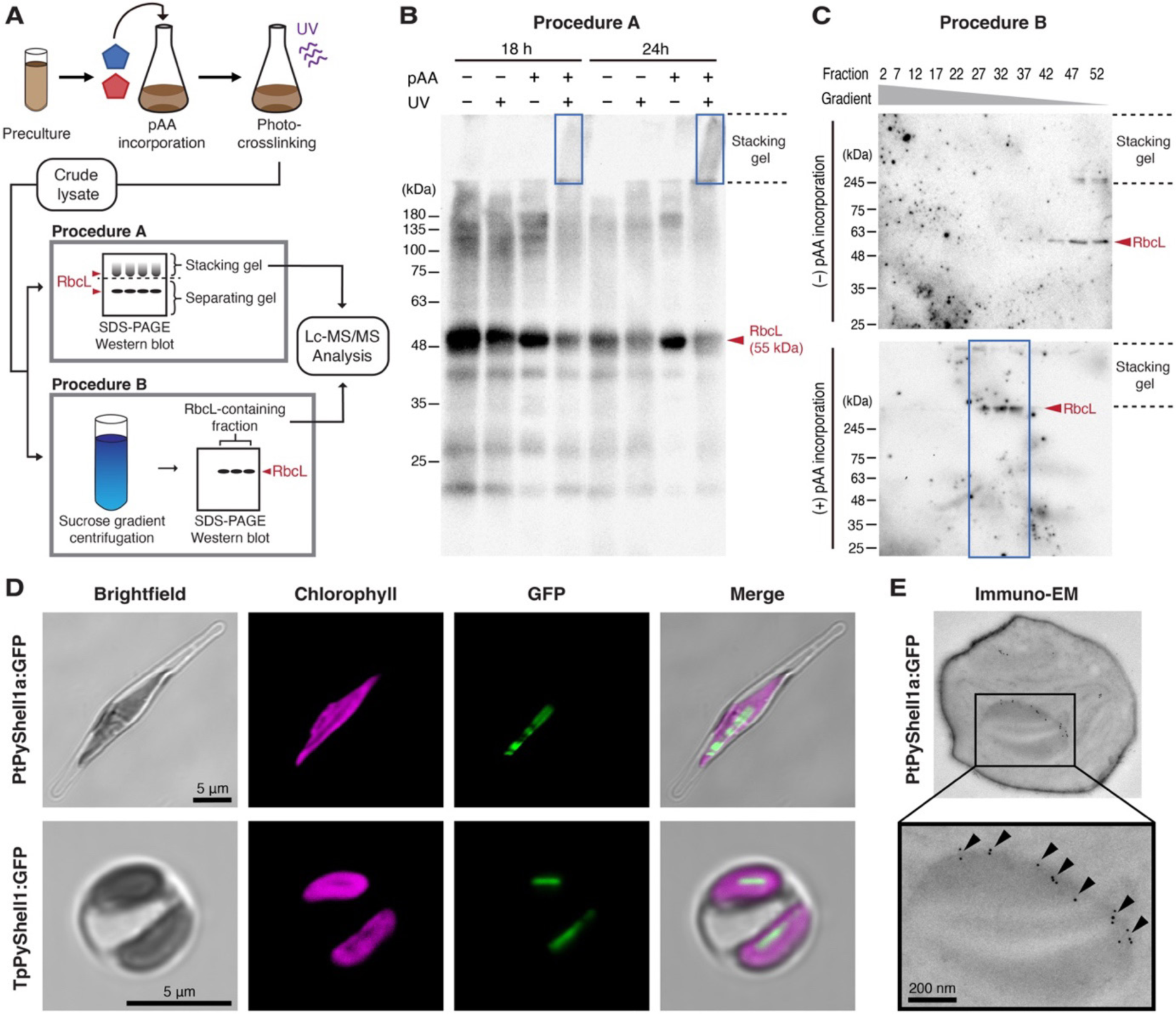
Identification of pyrenoid shell (PyShell) proteins in diatoms. **(A)** Proteomics-based workflow for detecting pyrenoid proteins in *P. tricornutum*. Cells were cultured with (+) or without (−) photo-reactive amino acids (pAA), photo-crosslinked *in vivo* with UV irradiation, and then disrupted by sonication. The crude extracts were subjected to either **(B)** SDS-PAGE (Procedure A) or **(C)** 22−55% (w/v) sucrose density gradient (Procedure B). Gel shift of the crosslinked Rubisco was detected by immunoblotting against the Rubisco large subunit (RbcL). Rubisco-containing gels or collected fractions (indicated by blue boxes in B and C) were digested by trypsin and analyzed by LC-MS/MS (for list of candidate Rubisco interactors, see Tables S1, S2). **(D)** Confocal images of PtPyShell1a:GFP in *P. tricornutum* (top row) and TpPyShell1:GFP in *T. pseudonana* (bottom row). See Fig. S2 for additional examples. **(E)** Immunogold-labeling TEM image of a *P. tricornutum* PtPyshell1a:GFP transformant probed with an anti-GFP antibody. Gold particles are indicated by black arrowheads. Scale bars: 5 µm in D; 200 nm in E.

From these two procedures, we identified more than 100 candidates for *P. tricornutum* pyrenoid proteins. We then filtered this list for the presence of the stramenopile-specific plastid targeting sequence (ER+ASAFAP) at the N-terminus (Gruber et al., 2007), yielding 49 and 14 candidate chloroplast proteins from Procedures A and B, respectively (Tables S1, S2). In addition to known pyrenoid proteins such as Rubisco, β-CAs (Tachibana et al., 2011), and FBAs (C1 and C5) (Allen et al., 2011), we identified several new candidates, which include an unknown protein (JGI protein ID: 45465), a putative acetyl-CoA carboxylase (JGI protein ID: 54926), cytochrome *c*_6_ (JGI protein ID: 44056), and a bestrophin-like protein (JGI protein ID: 46336). In the present study, we focused on the Pt45465 protein and its orthologues in the model diatoms *P. tricornutum* and *T. pseudonana*.

The *P. tricornutum* gene *Pt45465* (*PtPyShell1a*) is located on chromosome 7 together with a paralogue *Pt45466* (*PtPyShell2a*), which shares 74.3% similarity. There are duplications of both genes on chromosome 28: *Pt50215* (*PtPyShell1b*) and *Pt50214* (*PtPyShell2b*). All PtPyShell orthologs harbor stramenopile-type plastid targeting sequences. These proteins contain no transmembrane helices and thus are likely localized to the stroma. We defined two conserved regions (CR1 and CR2) in the PyShell proteins (Figure S1A), which were comprised of >50% hydrophobic amino acids. Using CR1 and CR2 as reference sequences, we searched for candidate Pyshell genes and found homologues primarily in diatoms and haptophytes, but also in a few marine algae from other clades (Figure S1C).

In *T. pseudonana*, we identified three putative PyShell orthologs: *Tp7881* (*TpPyShell1*), *Tp23918* (*TpPyShell2*), and *Tp7883* (*TpPyShell3*). Expression of these TpPyShell genes was analyzed in *T. pseudonana* wild-type cells grown under normal atmospheric CO_2_ (0.04%, hereafter “LC” for “low CO_2_”) and high CO_2_ (1%, hereafter “HC”), indicating that TpPyShell1 and 2 are the most abundant isoforms (Figure S1B).

We next checked the subcellular localization of PyShell proteins in *P. tricornutum* and *T. pseudonana* by fluorescence microscopy (Figures 1D, S2). We generated strains of these two diatom species expressing PtPyShell1a:GFP and TpPyShell1:GFP (C-terminal GFP tags), respectively. In *P. tricornutum*, PtPyShell1a:GFP signal was clearly detected in a hollow rod shape at the center of the chloroplast where the chlorophyll autofluorescence was dim, suggesting localization to the pyrenoid, perhaps surrounding the Rubisco matrix (Figures 1D, S2A-B). In *T. pseudonana*, we similarly observed TpPyShell1:GFP signal in a rod shape at the center of the chloroplast (Figures 1D, S2B). We further analyzed the *P. tricornutum* strain by immunoelectron microscopy, with anti-GFP nanogold localization confirming that the PtPyShell1a:GFP proteins were accumulated along the peripheral regions of the pyrenoid (Figure 1E).

### Molecular architecture of the PyShell lattice inside native diatom cells

To gain a higher resolution view of the pyrenoid periphery, we turned to *in situ* cryo-electron tomography (cryo-ET) (Turk and Baumeister, 2020; Hylton and Swulius, 2021). *P. tricornutum* and *T. pseudonana* cells were vitreously plunge-frozen on EM grids, thinned with a focused ion beam (Schaffer et al., 2017), and imaged in three dimensions with cryo-ET. We observed that the pyrenoids of both diatom species are surrounded by a proteinaceous sheath, which tightly encloses the Rubisco matrix (Figures 2A-D, S3). Closer inspection of these sheaths revealed that they are apparently formed from a repetitive lattice of protein subunits (Figures 2E-F; S3C, G). We hereafter refer to this pyrenoid-encapsulating shell as “the PyShell”; its location is consistent with our observations of PtPyShell1a:GFP and TpPyShell1:GFP, while our structural and mutational analysis described later in this study definitively implicate PyShell proteins in sheath formation.

**Figure 2.**
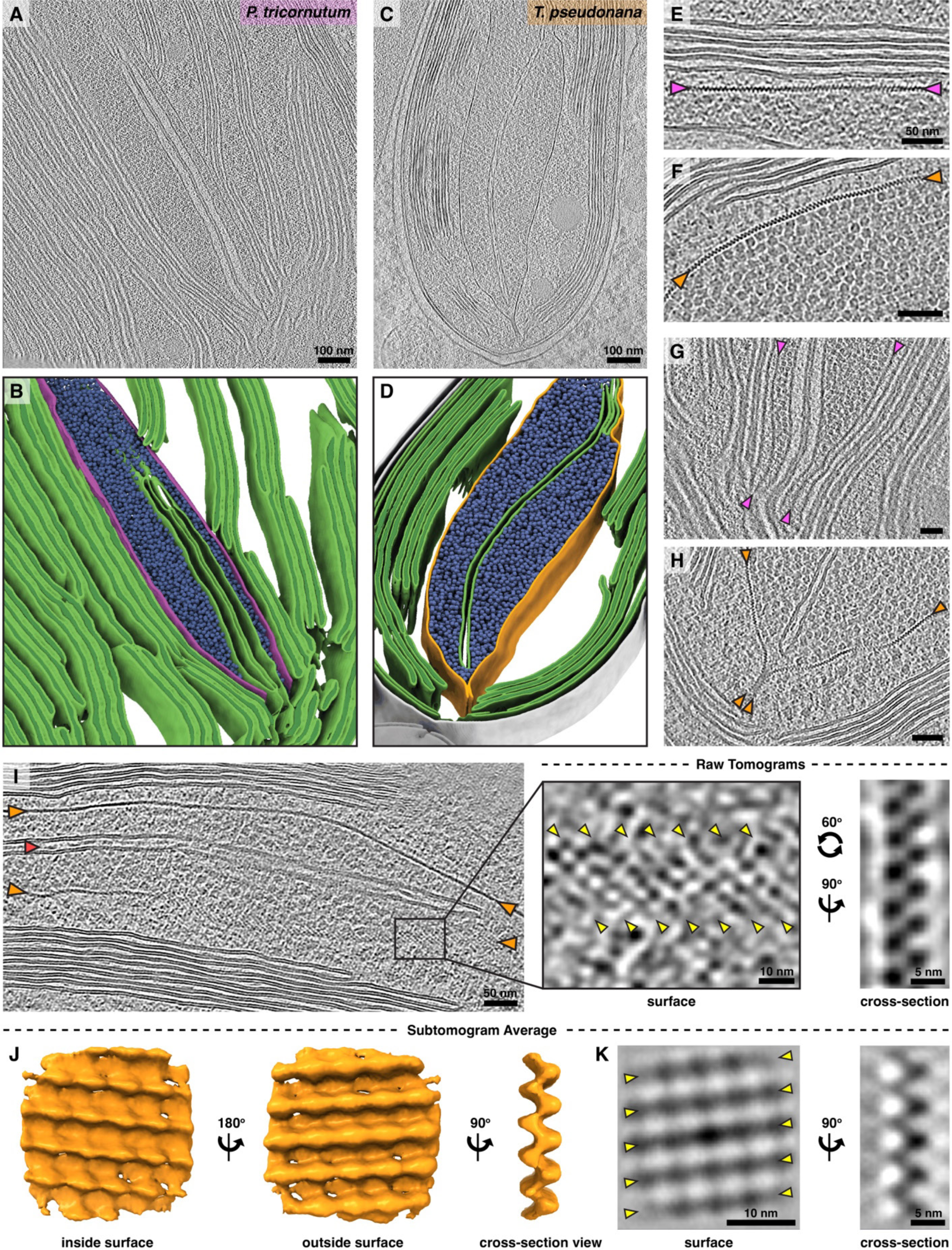
*In situ* cryo-ET reveals the native architecture of the PyShell inside diatom cells. Magenta labels and arrowheads: *P. tricornutum*. Orange labels and arrowheads: *T. pseudonana*. **(A,C)** 2D overview slices through tomograms and **(B,D)** corresponding 3D segmentations (green: thylakoids, blue: Rubisco complexes, magenta or orange: PyShell). **(E-F)** Close-up views of native PyShells (marked by arrowheads) in both diatom species. **(G-H)** Comparison of pyrenoid ends. In *P. tricornutum*, there is a gap in the PyShell that allows entry of two special thylakoids into the pyrenoid. In *T. pseudonana*, two apposing sheets of the Pyshell bind each other to seal the pyrenoid matrix. **(I)** Molecular details of the PyShell in raw tomograms. Left: overview revealing a stripe pattern when the PyShell twists to show its surface view (red arrowhead: particles inside the lumen of traversing thylakoid). Center: zoom in on the surface view, with the major stripes of the PyShell lattice marked with yellow arrowheads. Right: zoom in on a cross-section view, showing an apparent lattice of dimers. **(J-K)** Subtomogram average (STA) of the PyShell from *T. pseudonana*, shown in 3D isosurface view (J), as well as 2D slices (K) showing the surface view (yellow arrowheads: major stripes of lattice) and cross-section view. Scale bars: 100 nm in A-B; 50 nm in E-H and I, left; 10 nm in I, right and K, left; 5 nm in I, right and K, right. See Fig. S3 for additional cryo-ET images from both species.

The native cellular views provided by cryo-ET revealed some species-specific differences in pyrenoid architecture. In *P. tricornutum*, the PyShell is relatively flat and straight. Two specialized thylakoids penetrate the rubisco matrix and run the length of the pyrenoid (Figures 2A, G; S3B). The luminal space of these traversing thylakoids is swollen and sometimes filled with dense particles (red arrowheads). At the two ends of the pyrenoid, the PyShell closely associates with these two traversing thylakoids, which exit the pyrenoid and connect to the rest of the thylakoid network (Figures 2A, B, G; S3A). In *T. pseudonana*, the PyShell has more regions of high local curvature.

This pyrenoid is also bisected by one or two specialized thylakoids (Figures 2C, D, H, I; S3F) that frequently contain dense particles in their lumen; however, we never observed these thylakoids exiting the pyrenoid. Instead, at the two ends of the pyrenoid, the PyShell interacts with itself like a zipper to seal the Rubisco matrix (Fig 2C, D, H; S3E). Due to the limited cell area visualized by cryo-ET, we cannot definitely conclude that the pyrenoid thylakoids in *T. pseudonana* are disconnected from the rest of the thylakoid network. However, if such connection sites exist, they are much rarer than in *P. tricornutum.*

In our raw tomograms, the PyShell showed different features depending on its orientation. When observed in cross-section, it resembled a solid line, a chain of dots, or a zig-zag (Figure 2I, right). However, when the PyShell twisted 90 degrees to show its face, we could observe a lattice of subunits producing clear stripe patterns (Figure 2I, middle). To understand the three-dimensional structure of the PyShell, we performed subtomogram averaging (STA) (Wan and Briggs, 2016) of subvolumes picked along PyShell sheaths from our highest quality tomograms of *T. pseudonana*. After iterative alignment, classification, and polishing steps (see methods), we ultimately resolved a ∼20 Å density map of the native *T. pseudonana* PyShell using 14,341 subvolumes from seven tomograms (Figures 2J, S4). The STA density contains the stripe and zig-zag features seen in the raw tomograms, and reveals the 3D architecture of a tightly packed pseudocrystalline protein lattice.

### High-resolution *in vitro* structure of the *T. pseudonana* PyShell lattice

We required even higher resolution to determine precisely how individual PyShell proteins assemble to form a tight protein lattice. Therefore, we reconstituted the PyShell *in vitro* and performed single particle cryo-electron microscopy (cryo-EM). We expressed and purified recombinant TpPyShell1, which when concentrated *in vitro* to 2 mg/mL, self-assembled to form both flat sheets and hollow tubes with an outer diameter of 30 nm (Figure S5A-C). Following cryo-EM imaging and particle picking along the tubes, we used single particle analysis (SPA) and helical reconstruction to attain a 3.0-Å density map (Figures 3A, S5E-F), enabling us to build an atomic model of TpPyShell1 assembled in a homo-oligomeric lattice (Figure 3B-E, S5G).

**Figure 3.**
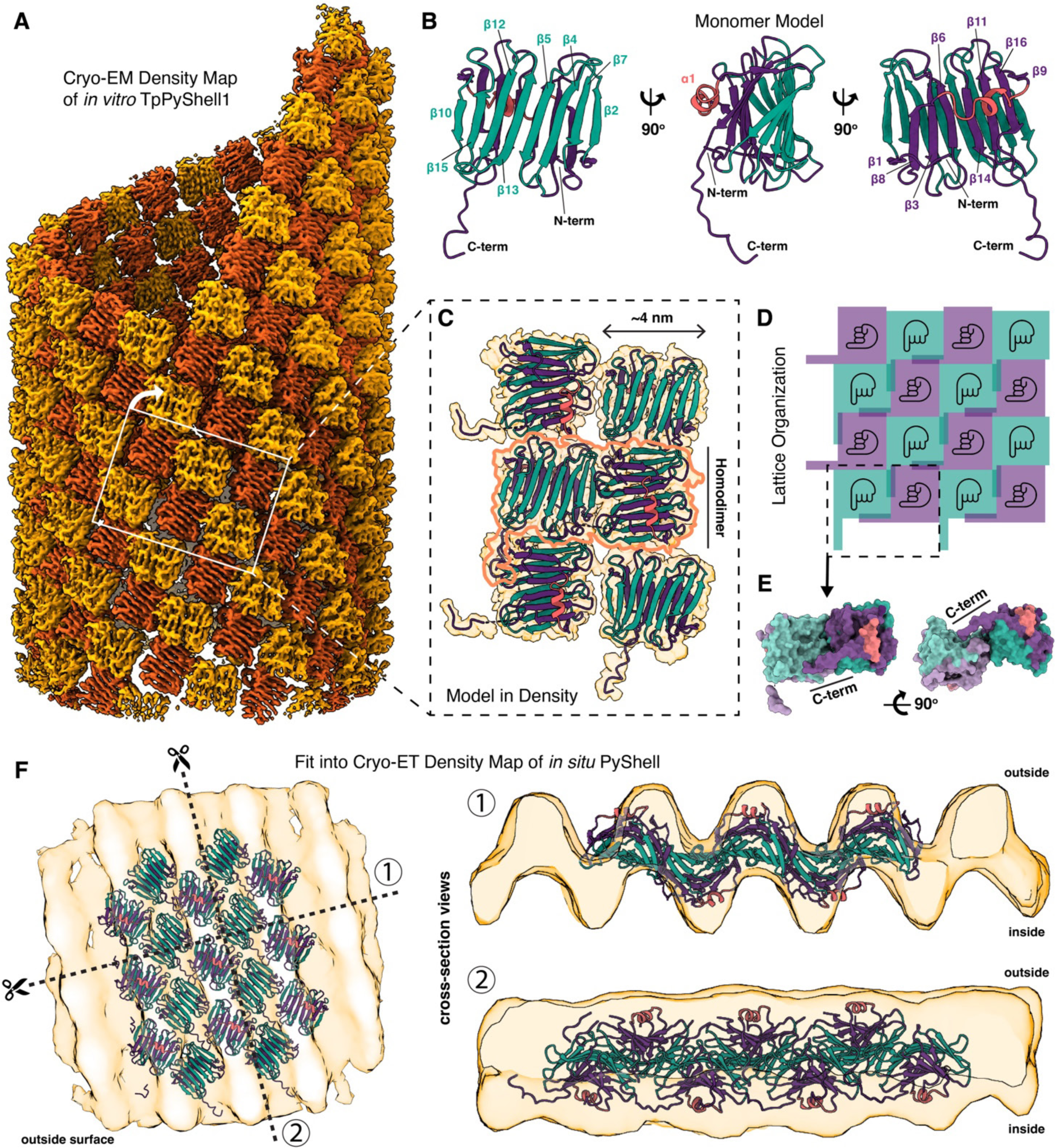
High-resolution *in vitro* structure of the *T. pseudonana* PyShell lattice. **(A)** Cryo-EM density map obtained by single particle analysis (SPA) and helical reconstruction of TpPyShell1, which assembles into a tube of *in vitro*. Global resolution: 3.0 Å (see Fig. S5E-F). **(B)** Cartoon model of the TpPyShell1 monomer. The two β-sheets (each composed of eight β-strands) and the adjacent α-helix are indicated in teal, purple, and pink, respectively. **(C)** Models of six TpPyShell1 monomers fit into the cryo-EM density map from A (yellow). The minimal building block of the tube’s lattice is a homodimer of TpPyShell1 proteins (outlined in orange), which are flipped and rotated 90 degrees relative to each other. **(D)** Schematic representation of this lattice arrangement, with hands indicating the flipping and rotation of monomers. The pinky finger represents the C-terminal domain (C-term). **(E)** Surface model representation of a homodimer unit from the lattice. The C-term extends and contacts a pocket in the adjacent monomer (also see Fig. S5J). **(F)** Fit of the unrolled SPA model (cartoon representation) into the *in situ* STA density from Fig. 2J (yellow). The surface view on the left is annotated with the cut directions corresponding to the two perpendicular cross-section views on the right (1 and 2).

The *in vitro* tube is an assembly of TpPyShell1 proteins in two alternating poses: half of the monomers face inward, while the other half face outward with a 90 degree in-plane rotation relative to the inward-facing monomers (Figure 3A, C-D). Thus, the minimal building block of the tube is a homodimer of TpPyShell1 proteins adopting these two poses (Figures 3E). The full tube has helical symmetry (rise of 25.14 Å, twist of −32.46°) and a symmetric unit consisting of seven homodimers (Figure S5G).

Each TpPyShell1 monomer contains 16 β-strands arranged in two parallel β-sheets, one slightly more extended than the other (Figure 3B, teal and purple for more and less extended sheets, respectively). The TpPyShell1 monomer has an internal pseudo-two-fold symmetry that subdivides the two β-sheets into the two conserved regions that we previously identified by bioinformatics: CR1 and CR2 (Figure S5H). A short α-helix spanning residues 169-181 (Figure 3B, S5H, pink) is positioned along the wall of the less extended β-sheet and connects CR1 with CR2. The N-terminal 68 residues were not well resolved in our density map and are likely flexible; a second short α-helix appears to be present in this region but could not be clearly modeled. The C-terminal domain extends from one monomer and contacts a positively-charged pocket on the adjacent monomer of the opposite pose, possibly providing a stabilizing interaction for the lattice (Figures 3E, S5J). We also observed small ∼1 nm gaps in the lattice at the junctions between four monomers; the residues surrounding these gaps do not have strong surface charge or hydrophobic properties (Figure S5I).

We next compared this high-resolution *in vitro* structure of a TpPyShell1 tube to our *in situ* structure of a PyShell sheet from inside *T. pseudonana* cells. To do so, we first unrolled the SPA density and fit the TpPyShell1 monomer models to form a flat lattice. It is noteworthy that, once flattened, the inside and outside surfaces of the *in vitro* TpPyShell1 lattice are practically identical and only distinguished by the spatial offset between alternating monomers. We next fit this flat lattice model into our *in situ* STA density map to produce a putative atomic model of how PyShell proteins may be arranged inside the cell (Figure 3F). The monomers of the SPA model matched the lattice spacing of the STA map (unrolling may had minor effects on the model spacing). Cross-sections through the fitted density (Figure 3F, right) reveal how the more extended β-sheets (teal) align to form a continuous tight wall at the center of the PyShell. The less extended β-sheets (purple) protrude inward and outward, positioning the short α-helixes (pink) on each lobe as the most distant domain from the central wall of the PyShell lattice. We note that there is some extra density at the tips of *in situ* lobes that extends further than the α-helices and is not occupied by the fitted *in vitro* TpPyShell1 lattice model.

### PyShell mutants have altered pyrenoid morphology and reduced CO_2_ fixation

To understand the physiological role of the PyShell *in vivo*, we performed simultaneous gene disruptions of *TpPyShell1* and *TpPyShell2* in *T. pseudonana* using a CRISPR/Cas9 (D10A) nickase approach that we recently developed for diatom gene editing (Nawaly et al., 2020). Because these two genes share high sequence similarity (92.8%), we were able to design a single set of guide RNAs targeting the CR1 domain of both genes (Figure S5A). Two independent biallelic double knock-out mutants were successfully obtained, denoted ΔTpPyShell1/2-1 (*m1*) and ΔTpPyShell1/2-2 (*m2*) (Figure S6A-B). Western blotting indicated that *m1* and *m2* lacked both the TpPyShell1 and TpPyShell2 proteins (Figure 4A).

**Figure 4.**
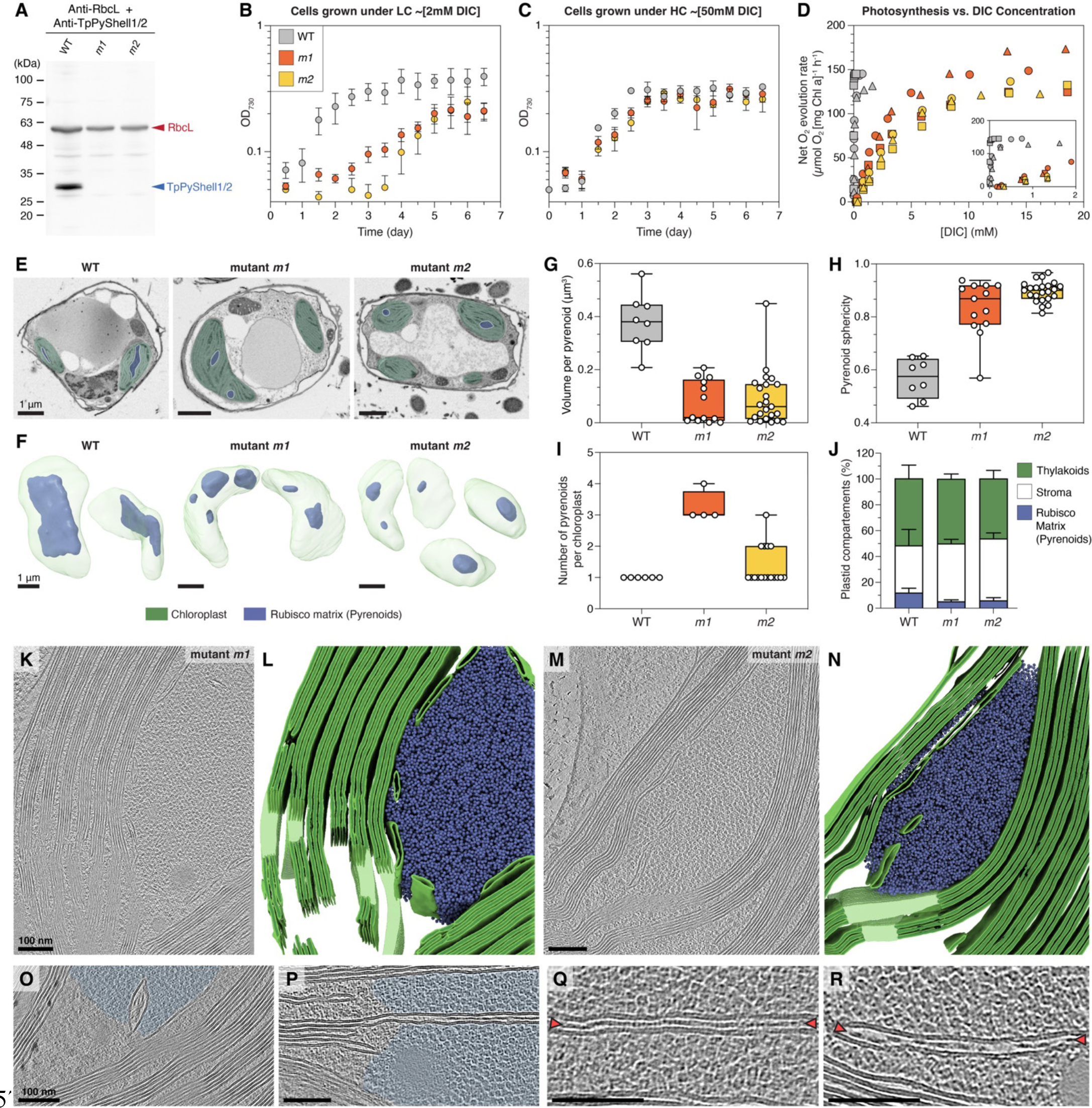
Phenotypes of *T. pseudonana* PyShell-deficient mutants. Mutants *m1* and *m2* (ΔTpPyShell1/2) have slightly different gene truncations (see Fig. S5A-C). **(A)** Western blot of crude cell extract (5 µg each) with anti-PyShell1/2 antibody, confirming the absence of TpPyShell1 and 2 proteins in the mutants **(B-C)** Growth of WT cells (grey), *m1* (orange), and *m2* (yellow) in air-level CO_2_ (LC; 0.04%; equivalent to ∼2 mM DIC) and high CO_2_ (HC; 1%; equivalent to ∼50 mM DIC) conditions, under continuous light (40 µmol photons m^−2^ s^−1^). Points: mean, error bars: standard deviation (*n* = 3, biological replicates). **(D)** Dependence of photosynthetic activity (measured by O_2_ evolution under 900 µmol photons m^−2^ s^−1^ constant actinic light) on DIC concentration (set by supplementing with bicarbonate) in WT cells (grey), *m1* (orange), and *m2* (yellow). Cells preconditioned in LC conditions. Data from three independent experiments are shown with different symbols. **(E-F)** FIB-SEM imaging of WT, *m1*, and *m2* cells, shown as (E) 2D slices through the raw tomographic data and (F) 3D segmentations of the chloroplast (green) and pyrenoid Rubisco matrix (blue) volumes. **(G-J)** Morphometric quantification of pyrenoids (Rubisco matrix regions) from FIB-SEM data: (G) volume per pyrenoid, (H) pyrenoid sphericity, (I) number of pyrenoids per cell, (J) percent chloroplast volume occupied by pyrenoid, thylakoids, and stroma. Box plots in G-I show median (center line), 75%-25% percentiles (box borders), and max-min values (whiskers). Error bars in J are standard deviation. N chloroplasts: 8 WT; 4 *m1*; 17 *m2*. **(K-R)** Cryo-ET of *m1* and *m2* cells. Overviews (K,M: tomographic slices, L,N: 3D segmentations) show higher sphericity of the Rubisco matrix and failure of specialized thylakoids to properly traverse the matrix. (O-P) Closeup tomographic slices showing the defined border of the Rubisco matrix (light blue) in the PyShell mutants. (Q-R) Closeup tomographic slices showing luminal particles (red arrowheads) inside the mislocalized specialized thylakoids. Scale bars: 1 µm in E-F, 100 nm in K-R. See Fig. S6 for additional cryo-ET images of the mutants.

Mutants *m1* and *m2* both showed severely inhibited growth in normal atmospheric CO_2_ (LC, “low CO_2_”) (Figure 4B). Compared to WT, the mutants had a longer lag phase, a slower growth rate, and took twice as long to reach stationary phase. In contrast, the growth profiles of WT, *m1*, and *m2* were similar when supplemented with 1% (HC, “high CO_2_”) (Figure 4C), This indicates that deletion of the major PyShell genes in *T. pseudonana* gives a clear high-CO_2_-requiring phenotype, presumably because of an impaired CCM. To test this hypothesis, we analyzed the photosynthetic affinity of WT and mutant cells for DIC by measuring net O_2_ evolution rate at increasing DIC concentrations (Figure 4D). Whereas WT cells reached their maximum photosynthetic rate (*P*_max_) at <0.5 mM [DIC], the mutants *m1* and *m2* required >10 mM DIC to reach their *P*_max_. Other photosynthetic parameters, including the DIC compensation point and apparent photosynthetic conductance, also support the highly impaired photosynthetic uptake of carbon in these mutants (Table S3), indicating that they cannot efficiently provide CO_2_ to Rubisco in seawater with less than 10 mM [DIC]. For perspective, the average [DIC] near the ocean surface is ∼2 mM (Cole et al., 2021).

To investigate how this high-CO_2_-requiring mutant phenotype is related to pyrenoid morphology, we reconstructed the 3D architecture of WT, *m1*, and *m2* cells using focused ion beam scanning electron microscopy (FIB-SEM). Cells were cryo-fixed at high-pressure to improve sample preservation, then subjected to freeze substitution and resin embedding. Following FIB-SEM imaging (Figure 4E), we used 3D segmentation to quantify the volumes and shapes of chloroplast regions (Figure 4F-J). In WT *T. pseudonana*, pyrenoids consisted of a single elongated compartment occupying around 10% of the chloroplast volume, similar our previous measurements of *P. tricornutum* (Uwizeye et al., 2021). Conversely, *m1* and *m2* chloroplasts contained multiple oval pyrenoid-like structures (higher sphericity than WT) of heterogeneous size, indicating that removal of the PyShell causes fragmentation of the Rubisco matrix and loss of its normal elongated architecture. Furthermore, the total measured volume of these mutant pyrenoids decreased to <5% of the chloroplast in the mutants (Figure 4J), suggesting that a portion of the Rubisco may be disperse in the stroma or present as small aggregates not detectable by our FIB-SEM imaging.

We next performed *in situ* cryo-ET of *m1* and *m2* cells to gain a molecular-resolution view of the mutant pyrenoids (Figure 4K-R). Consistent with the FIB-SEM imaging, we observed pyrenoid-like aggregates of Rubisco matrix that were more oval in shape than WT pyrenoids. With the resolution afforded by cryo-ET, we confirmed that these mutant pyrenoids were not encased in a PyShell sheath. Nevertheless, the cohesion of the Rubisco matrix was maintained without the PyShell, consistent with a recent report of a linker protein that mediates phase separation of Rubisco in diatoms (Oh et al., 2023). The rounder shape of the mutant pyrenoids and clear boundary between the Rubisco matrix and the stroma (Figure 4O-P, S6E) are also consistent with phase separation. Strikingly, the specialized thylakoids with luminal particles that normally traverse the long axis of WT pyrenoids were strongly disturbed in the mutants. Thylakoids containing luminal particles were frequently seen in the peripheral regions of mutant pyrenoids (Figure 4Q-R, S6E) but often appeared fragmented and did not pass through the matrix center (Figure 4K-N). Therefore, the PyShell appears to play important roles both in maintaining the elongated shape and singular cohesiveness of *T. pseudonana* pyrenoids, as well as helping position the specialized thylakoids on an end-to-end trajectory bisecting the Rubisco matrix.

## Discussion

In this study, we discovered the PyShell, a protein sheath that encases the pyrenoids of *P. tricornutum* and *T. pseudonana*, model species of the pennate and centric diatom clades, respectively (Figure 1). We characterized the structure of the *T. pseudonana* PyShell lattice across scales, from near-atomic resolution *in vitro* to molecular resolution within native diatom cells (Figures 2-3). Our functional analysis of *T. pseudonana* PyShell deletion mutants showed that the PyShell sheath maintains pyrenoid architecture and is essential for efficient function of the CCM, and thereby the ability of diatoms to grow by assimilating environmental CO_2_ (Figure 4).

How does the PyShell contribute so significantly to diatom CCM function? Comparison to the most well-characterized pyrenoid system in the green alga *C. reinhardtii* may help provide some insight. Reaction-diffusion modeling of *C. reinhardtii* (Fei et al., 2022) suggests that all pyrenoid-based CCMs require the following essential features: 1) an aggregation of most of the chloroplast’s Rubisco enzymes in one location, 2) a local source of high CO_2_ concentration at the center of this Rubisco aggregate, and 3) a diffusion barrier at the aggregate border to prevent CO_2_ leakage. Our data indicate that the PyShell contributes to the first two essential pyrenoid features, and we speculate that the PyShell may directly perform the third (Figure 5).

**Figure 5.**
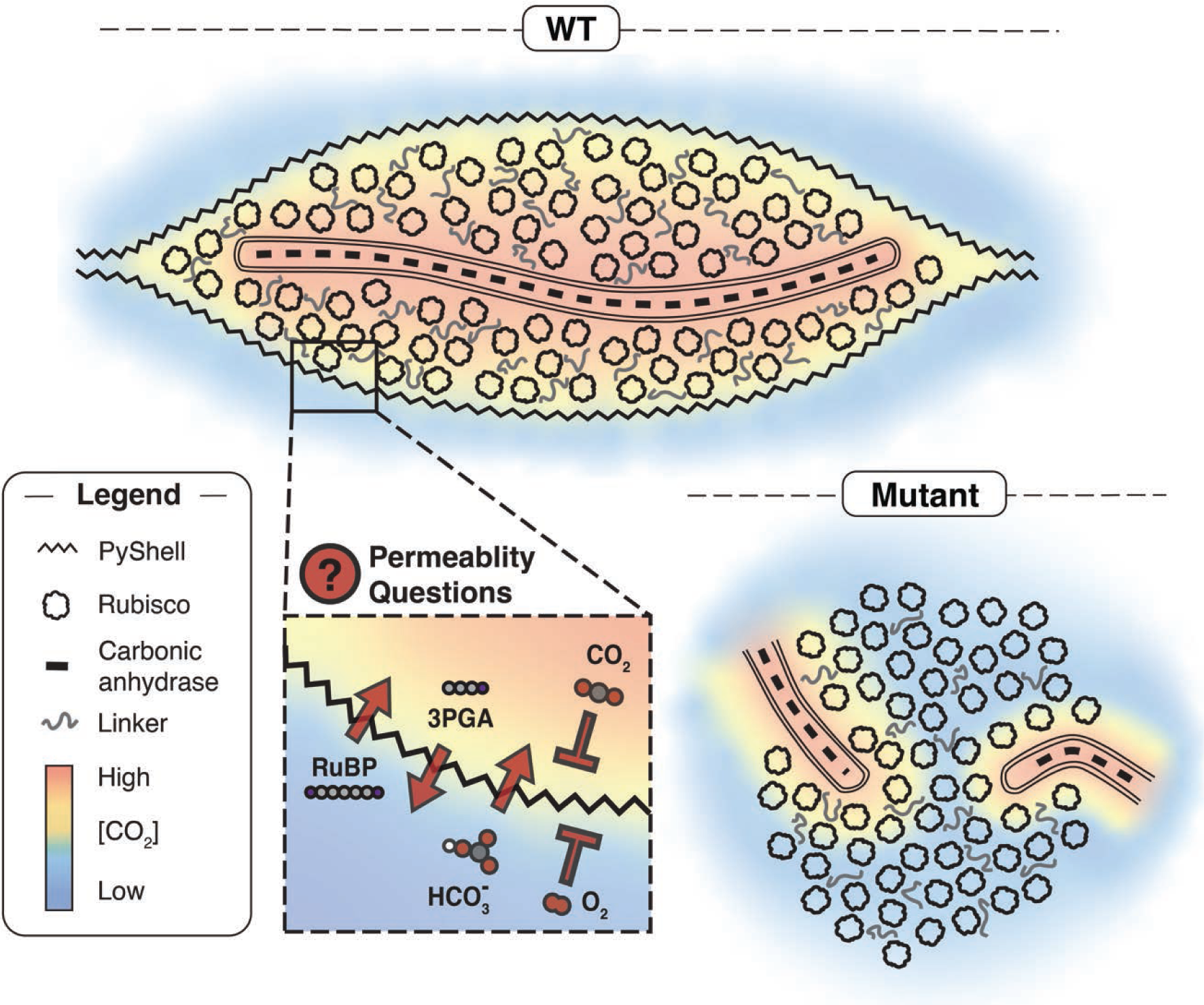
Proposed model of Pyshell funtion and open questions. In wild-type cells (WT), the PyShell encloses the Rubisco matrix, enforcing an elobngated pyrenoid shape. One or two specialized thylakoids traverse the long axix of the pyrenoid, Carbonic anhydrase inside the lumen of the these thylakoids generates Co_2_, which diffuses throught the thylakoid membrances and permeates the Rubisco matrix, enabling efficient carbon fixation. It remains to be determined how bicarbonate (hCO3) reaches the luminal carbonic anhydrase,and how sugar substrates (Ribulose 1,5-bisphosphate; RuBP) and products (3-Phosphoglyceric acid; 3PGA) transit between the stroma and the Pyshell-enveloped Rubisco matrix. We speculate that the PyShell may also serve as a gas diffusion barrier, keeping O_2_ out of the pyrenoid while maintaining the high local concentration of CO_2_. In the PyShell-deficient mutants (Fig. 4), the Rubisco matrix remains aggregated by a linker protein but forms a more spherical shape. The specialized thylakoids fail to bisect the matrix, delocalizing the source of CO_2_ from the pyrenoid center. If the PyShell fundtions as an O_2_/CO_2_ diffustion barrier, its absence would further inhibit Rubisco activity. This defective CCM underlines the mutants’ high-Co_2_ requiring phenotype.

In *C. reinhardtii*, the molecular mechanism underlying aggregation of pyrenoid Rubisco (essential feature #1) has been well characterized. The multivalent linker protein EPYC1 is necessary and sufficient for Rubisco to undergo liquid-liquid phase separation both *in vitro* and *in vivo*, giving rise to a condensed yet fluid pyrenoid matrix (Mackinder et al., 2016; Freeman Rosenzweig et al., 2017; Wunder et al., 2018; He et al., 2020). Rubisco phase separation appears to be common throughout evolution, as Rubisco has been observed to form liquid condensates during the biogenesis of cyanobacterial carboxysomes (Wang et al., 2019; Oltrogge et al., 2020; Zang et al., 2021). Evidence is now mounting that the Rubisco matrix of diatom pyrenoids also forms by phase separation. Recently, the multivalent linker protein PYCO1 was shown to localize to the *P. tricornutum* pyrenoid matrix and phase separate with Rubisco *in vitro* (Oh et al., 2023). Furthermore, our FIB-SEM and *in situ* cryo-ET of the *T. pseudonana* ΔTpPyShell1/2 mutants (*m1* and *m2*) show that Rubisco continues to aggregate in the absence of the PyShell, forming rounder bodies with clear boundaries between Rubisco and stroma that are consistent with phase-separated condensates (Figures 4E,F, H, K-P; S6E). The PyShell is thus not required for phase separation of Rubisco. However, it may still play a role in concentrating the majority of Rubisco at one spot. In the *m1* and *m2* mutants, Rubisco forms multiple smaller condensates throughout the chloroplast, and the combined volume of these dispersed condensates is less than a single WT pyrenoid, suggesting that some Rubisco may not enter the condensed phase (Figure 4E-G, I). We speculate that a direct or indirect interaction between the PyShell and Rubisco helps concentrate all the condensed Rubisco at a single location in the chloroplast. Indeed, in some cryo-ET images, it appears that a single layer of ordered Rubisco is lined up along the PyShell lattice (Figures 2F; S3D,H). The precise mechanism of this direct or indirect PyShell-Rubisco interaction requires further investigation.

*C. reinhardtii* and diatoms use a common strategy to produce a local source of high CO_2_ concentration at the center of the pyrenoid’s Rubisco matrix (essential feature #2). Both algae have thylakoid-derived membrane systems that cross the center of the pyrenoids, taking the shape of cylindrical tubules in *C. reinhardtii* (Ohad et al., 1967; Engel et al., 2015) and specialized thylakoid sheets in *P. tricornutum* and *T. pseudonana* (Pyszniak and Gibbs, 1992; Bedoshvili et al., 2009). Inside the luminal space of these pyrenoid-traversing tubules and thylakoids are carbonic anhydrases (CAs; α-type CAH3 in *C. reinhardtii,* θ-type Ptθ-CA1 in *P. tricornutum* and Tpθ-CA2 in *T. pseudonana*) (Karlsson et al., 1998; Kikutani et al., 2016; Nawaly et al., 2023; Shimakawa et al., 2023), which convert HCO_3_^−^ into CO_2_ at the center of the pyrenoid. This helps maximize Rubisco efficiency by saturating the enzyme with its CO_2_ substrate and outcompeting unproductive binding of Rubisco to O_2_. However, in the *T. pseudonana m1* and *m2* mutants, the specialized thylakoids are mislocalized to the periphery of the Rubisco condensate, and therefore, cannot provide a source of CO_2_ at the center of the pyrenoid (Figure 5). Our cryo-ET observations clearly show that the PyShell is necessary to correctly position these specialized thylakoids along the long axis of the Rubisco matrix (Figure 4K-N). By constricting the pyrenoid matrix into a long tube surrounding the CA-containing thylakoids, the PyShell minimizes the diffusion distance of CO_2_ from CA-source to Rubisco-sink.

Modeling indicates that CCM efficiency is greatly enhanced when a pyrenoid’s Rubisco matrix is surrounded by a diffusion barrier to limit CO_2_ leakage (essential feature #3). In *C. reinhardtii*, the identity of this barrier is debated, with candidates including a pyrenoid-peripheral CA and the pyrenoid’s surrounding starch sheath (Ramazanov et al., 1994; Fei et al., 2022). In diatoms, the PyShell forms a tight sheath around the Rubisco matrix that larger molecules certainly cannot pass. In this way, the PyShell is analogous to the proteinaceous shells of cyanobacterial carboxysomes (Shively et al., 1973; Melnicki et al., 2021), which similarly form a dense wall around an aggregate of encapsulated Rubisco. There is experimental evidence that carboxysome shells may be selectively permeable, blocking diffusion of CO_2_ and O_2_, while permitting passage of HCO_3_^-^ through small pores in the wall (Dou et al., 2008; Cai et al., 2009; Mahinthichaichan et al., 2018; Faulkner et al., 2020; Huang et al., 2022). In this way, HCO_3_^-^ would diffuse into the carboxysome, where it is converted to CO_2_ by a CA, trapping a high concentration of CO_2_ with Rubisco inside the carboxysome shell. Similarly, there is evidence that carboxysome shell pores can be gated with an airlock-like mechanism to allow selective passage of sugar metabolite substrates and products of Rubisco CO_2_ fixation (Klein et al., 2009; Cai et al., 2013; Larsson et al., 2017). Our high-resolution *in vitro* structure of a homo-oligomerized TpPyshell1 tube revealed ∼1 nm gaps in the otherwise densely packed lattice (Figures 3C, S5I). However, to understand whether the PyShell functions as a selective barrier, detailed investigations are required to probe the PyShell’s permeability to small molecules, in particular CO_2_, O_2_ and the sugar metabolites RuBP and 3PGA (Figure 5, permeability questions).

In summary, the *m1* and *m2* PyShell deletion mutants exhibit such a strong inhibition of carbon fixation efficiency and growth (Figure 4B-D) because they may have defects in all three essential features of pyrenoid CCMs: Rubisco aggregation at one location of the chloroplast, a CO_2_ source at the center of the Rubisco aggregate, and a barrier to prevent CO_2_ leakage. Our physiological measurements (Figures 4D, S6C-D) show that the PyShell-deficient mutants require ∼10 mM DIC to saturate their photosynthesis, which is roughly 5-fold higher than the concentration of DIC in the ocean. Thus, the PyShell is likely essential to maintain efficient carbon fixation of diatoms in the wild.

Our bioinformatic analysis indicates that PyShell orthologs are widespread mainly in diatoms and haptophytes (Figure S1C). Although these two clades are not closely related phylogenetically, both have a plastid of red algal origin. Orthologs are also found some non-diatom stramenopiles, including pelagophytes and dictyochophytes, as well as several species of alveolata. It is plausible that the PyShell originated from the photosynthetic ancestor of haptophytes and the SAR supergroup (stramenopiles, alveolates, Rhizaria), and thereafter evolved independently in each clade. To understand the global prevalence of the PyShell, we queried the Ocean Gene Atlas (Villar et al., 2018) and found that PyShell transcripts are broadly detected in marine environments around the world (Figure S1D), with almost all these sequences belonging to stramenopiles and haptophytes. These abundant taxa produce immense biomass through the photosynthetic uptake of CO_2_, which provides an energy source for much of the life in the ocean. The PyShell therefore plays a major role in the CCM of environmentally-relevant marine algae, which account for roughly half of carbon fixation in the ocean, and by extension, one quarter of the carbon fixation on our planet. To forecast the future of the global carbon cycle, it will be important to understand how well PyShell-mediated carbon fixation can adapt to rapidly accelerating climate change. Major efforts are underway to engineer cyanobacterial carboxysomes and algal pyrenoids into plants to increase their carbon-fixation capacity (Hennacy and Jonikas, 2020; Borden and Savage, 2021). It is estimated that introducing such a CCM could increase yield by up to 60%, while reducing water and fertilizer requirements (McGrath and Long, 2014). Recent progress has been made in assembling components of the *C. reinhardtii* pyrenoid inside *Arabidopsis thaliana*, including the EPYC1 linker, which causes Rubisco to condense to form a pyrenoid-like matrix within these plant chloroplasts (Atkinson et al., 2020). Our discovery and characterization of the diatom PyShell expands the molecular toolbox with a lattice-forming protein that may be able to encapsulate these Rubisco condensates, providing a boundary between the artificial pyrenoid and the surrounding chloroplast. These engineering efforts hold potential for designing crops that grow faster, consume less resources, and are more resistant to environmental stress, helping feed the world’s growing population in regions of the planet that climate change is rapidly making less arable. Improving biological carbon capture may even one day help mitigate climate change itself, removing more CO_2_ from the atmosphere to slow the course of global warming.

### Limitations of the Study

Our study describes the PyShell in two model diatom species and demonstrates how this Rubisco-encasing protein barrier is essential for pyrenoid architecture and function in *T. pseudonana*. However, our data has some limitations that raise new questions requiring future in-depth investigation:

1. <u>PyShell structure</u>: The N-terminal 68 residues in TpPyShell1 are mostly disordered and were not resolved in our *in vitro* SPA structure. When mapping the unrolled SPA model into our *in situ* STA density (Figure 3F), we noted extra density beyond each monomer’s single α-helix. STA would require sub-nm resolution to distinguish whether this extra density is constituted by the PyShell lattice, itself, or rather comes from an interacting protein. This ambitious goal faces the challenges of resolving small proteins *in situ* (Russo et al., 2022), as well as potential heterogeneity of the PyShell within the cell. Mutagenesis studies of this α-helix may also help elucidate whether it mediates specific interactions. The biggest mystery from our SPA data is that the inside and outside surfaces of the TpPyShell1 lattice are structurally interchangeable (Figure 3F). Thus, additional studies are required to understand whether the two surfaces of the PyShell are distinct *in vivo*. For example, in the pyrenoid, Rubisco is only bound to the PyShell’s inner surface, but we cannot say from our data whether this is driven by asymmetry of the PyShell surfaces or rather is a consequence of pyrenoid biogenesis events.
2. <u>PyShell heterogeneity</u>: There are multiple homologous PyShell genes in both *P. tricornutum* and *T. pseudonana* (Figure S1A), with the latter expressing both TpPyShell1 and TpPyShell2 at high levels *in vivo* (Figure S1B). To resolve a high-resolution PyShell structure, we assembled the *in vitro* tube from only TpPyShell1, which also proves that a homogenous solution of this single protein is sufficient to assemble a lattice. It is quite possible that multiple PyShell isoforms hetero-oligomerize to make the PyShell lattice *in situ*. However, cryo-ET lacks the resolution to distinguish this heterogeneity due to the high homology and predicted structural similarity of different PyShell proteins. Complementary approaches combining mutagenesis and fluorescent tagging of each protein will be required to address whether PyShell isoforms co-assemble *in vivo*, whether there is a functional significance to the different isoforms, and whether these isoforms have discrete localizations within the pyrenoid sheath.
3. <u>PyShell permeability</u>: Our *in vitro* SPA structure shows small ∼1 nm gaps at the junctions between four monomers in the TpPyShell1 lattice (Figures 3C, S5I). While these gaps are potentially large enough to allow gas and some small metabolites to pass through the PyShell wall, in-depth functional studies would be required to assay PyShell permeability. These studies must take into consideration that the *in vivo* PyShell may be heterogeneous and also bound by interacting proteins.
4. <u>Pyrenoid architecture and biogenesis</u>: Our cryo-ET data provides a detailed look at the molecular architecture of steady-state pyrenoids in *P. tricornutum* and *T. pseudonana*, but it lacks insights into how this architecture is established. With the Rubisco matrix apparently completely encapsulated (except the thylakoid entry points in *P. tricornutum*), how does the pyrenoid grow during the cell cycle to incorporate new Rubisco and PyShell components? In both species, we find dense particles within the lumen of pyrenoid-traversing thylakoids (Figures 2I; 4Q-R; S3B,F). This is the location thought to be occupied by a carbonic anhydrase, but we cannot assign an identity to these luminal particles from cryo-ET alone. It also remains unanswered why these thylakoids acquire the luminal particles but fail to properly traverse the Rubisco matrix the ΔTpPyShell1/2 mutants.
5. <u>PyShell prevalence and architectural diversity</u>: Our bioinformatic analysis indicates that PyShell homologues are commonly found throughout diatoms and haptophytes, and they may also be present in other algal clades (Figure S1C). However, this analysis is biased towards sequenced species, so some clades may be underrepresented. Even in our cryo-ET comparison between two diatom species, we noted substantial differences in PyShell architecture, in particular at the pyrenoid ends (Figures 2A-D, G-H; S3A,E). Capturing the variations in PyShell architecture across evolution will require extensive cryo-ET of diverse algae, including in non-model species sampled directly from the environment (Mocaer et al., 2023).

## Materials and methods

### Cultures

The marine diatoms *P. tricornutum* Bohlin (UTEX642) and *T. pseudonana* (Hustedt) Hasle et Heimdal (CCMP 1335) were axenically and photoautotrophically cultured in artificial seawater medium with the addition of half-strength Guillard’s ‘F’ solution (Guillard and Ryther, 1962; Guillard, 1975) supplemented with 10 nM sodium selenite under continuous light (20°C, 40 μmol photons m^−2^ s^−1^, fluorescent lamp). The cultures were aerated with ambient air (0.04% CO_2_) or 1% CO_2_ gas for LC or HC conditions, respectively. For the culture of *T. pseudonana*, the concentration of NaCl was lowered to 270 mM in the medium.

### *In vivo* cross-linking with photo-reactive amino acids

*P. tricornutum* cells grown in LC were harvested at logarithmic growth phase and resuspended in fresh medium at a concentration of OD_730_ = 1.0−1.2, in the presence of 1 mM ʟ-photo-leucine and 2 mM ʟ-photo-methionine (Thermo Fisher Scientific, Waltham, MA, USA). Incorporation of these photo-reactive amino acids (pAA) was performed under illumination with blue (455 nm) and red (635 nm) LED light (50 μmol photons m^−2^ s^−1^) for 6−24 hours. Subsequently, the cell cultures were irradiated with UV light (365 nm) for 30−45 min to perform *in vivo* photo-cross-linking. Cells were harvested and resuspended in 25 mM Tris-HCl (pH 7.0), then disrupted by sonication. Insoluble debris was removed by centrifugation, and the resulting supernatant was either subjected to SDS-PAGE (for “gel digestion, procedure A”) or centrifuged on a 25−55% (w/v) linear sucrose gradient in 25 mM Tris-HCl (pH 7.0) at 210,000 × *g* for 4 h at 4°C (for “solution digestion, procedure B”). Aliquots (200 μL) of each fraction were collected, and the protein concentrations were determined with a protein assay kit (Bio-Rad, Hercules, CA, USA) using bovine serum albumin as a standard.

### Western blotting

Proteins extracted as described above were electrophoretically separated by SDS-PAGE, transferred to PVDF membrane, and blocked with 1% (w/v) skim milk dissolved in phosphate-buffered saline (PBS) containing 0.05% (v/v) Tween 20. For detection of RbcL, a rabbit anti-RbcL antiserum generated against *P. tricornutum* RbcL partial peptide (Japan Bio Serum, Hiroshima, Japan) was used as the primary antibody (diluted 1:1000). Goat anti-rabbit IgG conjugated with horseradish peroxidase was used as the secondary antibody (diluted 1:10000). Immunoreactive signals were detected by an enhanced chemiluminescence reagent (ImmunoStar Zeta, Wako, Osaka, Japan) with a high sensitivity CCD imaging system (Luminograph I, ATTO, Tokyo, Japan).

For the confirmation of the deletions of TpPyShell1 and 2 in *m1* and *m2*, we disrupted the *T. pseudonana* WT and mutants grown under HC by sonication in 50 mM HEPES (pH 7.5) with a protease inhibitor cocktail (nacalai, Kyoto, Japan) to obtain the crude extracts. Each 5 µg of protein were loaded and analyzed by immunoblotting as mentioned above. A rabbit anti-TpPyShell1 and 2 antiserum (Japan Bio Serum) targeting the conserved peptide sequence “GTARDLAEIWDNSS” was used as the primary antibody. The anti-RbcL antibody was also used as a loading control.

### Identification of proteins by LC-MS/MS

Proteins either in acrylamide gel (“Procedure A”) or solution (“Procedure B”) were subjected to reduction, alkylation, and digested by trypsin before injection into the LC-MS/MS. For gel samples, gel blocks (ca. 1 mm^3^) were dehydrated in 100 μL acetonitrile for 10 min at room temperature. After the removal of acetonitrile, the gel block was dried in an evaporator and incubated in 25 mM NH_4_HCO_3_ containing 10 mM dithiothreitol for 1 h at 56°C. The gel block was washed in 100 μL of 25 mM NH_4_HCO_3_ and incubated with 55 mM iodoacetamide for 45 min at room temperature. After washing twice with 100 μL of 25 mM of NH_4_HCO_3_, the gel block was dehydrated in acetonitrile. The dried gel blocks were soaked in 50 mM NH_4_HCO_3_ containing 10 ng μL^−1^ trypsin at 37°C for 16−20 h. Digested peptides were extracted with 50% (v/v) acetonitrile containing 5% (v/v) formic acid, concentrated by an evaporator, and dissolved in 1% (v/v) formic acid. The solutions were desalted by ZipTip C18 (Merck Millipore). For solution samples, disulfide bonds were reduced in 50 mM Tris-HCl (pH 8.5) containing 10 mM dithiothreitol at 37°C for 1.5 h, and subsequently alkylated with 50 mM iodoacetamide for 30 min at room temperature. Proteins were digested with 2 ng µL^−1^ trypsin in 50 mM NH_4_HCO_3_ for 16−20 h at 37°C, and then the reaction was stopped by addition of 0.18% (v/v) formic acid. Peptide samples were concentrated in an evaporator, dissolved in 0.1% (v/v) formic acid, and then desalted by ZipTip C18 (Merck Millipore, Burlington, MA, USA). The digested samples were injected into EASY-nLC 1000 connected to LTQ Orbitrap XL (Thermo Fisher Scientific). Data of LC-MS/MS were analyzed by the software Proteome Discoverer 1.4 (Thermo Fisher Scientific) with the open genome data resource for *P. tricornutum* from JGI (Phatr2).

### Genome editing of *TpPyShell1/2*

A pair of single guide RNA (sgRNA) sequences were constructed to target both *TpPyShell1* and *TpPyShell2* using ZiFiT Targeter (http://zifit.partners.org/ZiFiT/) (Sander et al., 2010), followed by Cas-Designer (http://www.rgenome.net/cas-designer/), which considers specificity and microhomology-mediated joining (Park et al., 2015). The nucleotides 5ʹ-TAACGGCATTGAAGGTACGA-3ʹ (396–415 in *TpPyShell1* and 423–442 in *TpPyShell2*) and 5ʹ-TCCCCGCGGCCCCAACACCG-3ʹ (393–412 in *TpPyShell1* and 420–439 in *TpPyShell2*) were chosen (Figure S6A). For appropriate RNA transcription, an additional G was inserted at the 5ʹ end of the DNA sequence encoding each sgRNA (Sander and Joung, 2014). The dual sgRNA vector targeting *TpPyShell1/2* was generated and introduced together with Cas9 (D10A) nickase vector into the *T. pseudonana* cells by particle bombardment according to the previous work (Nawaly et al., 2020). The mutant strains were screened on agar medium supplemented with 100 µg/mL nourseothricin (Jena Bioscience) under HC conditions. To ensure monoclonal strains, each colony was restreaked four times, each time picking a single colony from the streak. Mutations were verified by TIDE analysis and direct sequencing (Figure S6B).

### qRT-PCR of PyShell Transcripts

Transcript levels of *TpPyShell1*, *TpPyShell2*, *TpPyShell3*, and *TpδCA3* (Tp233) were quantified by qRT-PCR. *TpδCA3* was used as a control low-CO_2_ inducible gene. The internal controls were *Actin* (*Tp25772*), *GapC3* (*Tp28241*), and *Histone H4* (*Tp3184*), which were confirmed to be unresponsive to different CO_2_ concentrations in *T. pseudonana* (Matsui et al., 2018).Transcript levels were calculated with the 2^−ΔΔCt^ method against each internal control separately (Livak and Schmittgen, 2001). Then, the average values of ΔΔCt were calculated for each replicate.

### Expression of GFP fusion proteins in *P. tricornutum* and *T. pseudonana*

Correct full-length coding sequences for PyShell orthologues in *P. tricornutum* and *T. pseudonana* (PtPyShell1, TpPyShell1, and TpPyShell2) were determined by RACE using a SMARTer RACE 5’/3’ kit (TaKaRa). Sequences were amplified by PCR and cloned into pPha-T1 or pTha-NR vectors containing a fragment of enhanced GFP by a seamless ligation cloning extract method (Motohashi, 2015). The resulting plasmids were introduced into each WT cell using particle bombardment (PDS-1000/He, BioRad, Tokyo, Japan), and transformants expressing GFP were screened by fluorescence microscopy (Tachibana et al., 2011). Primers used are listed in Table S4.

### Confocal fluorescence microscopy

To observe subcellular localizations of GFP fusion proteins, we used confocal laser microscopes A1 (Nikon, Tokyo, Japan) or SP8 (Leica, Wetzlar, Germany). When imaging with the A1, chlorophyll autofluorescence was detected at 662–737 nm after excitation with a 638 nm laser, and GFP fluorescence was monitored at 500–550 nm following excitation at 488 nm. When imaging with the SP8, chlorophyll was excited by a 552 nm laser and detected at 600–750 nm. GFP was excited by a 488 nm laser and detected at 500–520 nm.

### Immunoelectron microscopy

The strain of *P. tricornutum* that expressing PtPyShell1 tagged with GFP was fixed by as previously described (Kikutani et al., 2016) with small modifications in the polymerization step; the samples immersed in resin were polymerized at −30°C for 5 days under UV light. Thin sections cut with Leica EM UC7 were mounted on nickel slot grids, followed by an edging step with 1% (w/v) sodium periodate. After the blocking step, the sections were reacted with polyclonal anti-GFP antibody (AnaSpec, Fremont, CA, US) diluted 1:500 in 3% (w/v) BSA in PBS at 25°C overnight. After rinsing with PBS, they were incubated for 60 min at room temperature with a goat anti-rabbit IgG conjugated to 10-nm colloidal gold particles (1:50 diluted in PBS; BBI Solutions, Crumlin, UK). The thin sections were stained with TI blue (Nisshin EM, Aichi, Japan), following washing with distilled water. The sections were observed with a JEM-1011 electron microscope (JEOL, Tokyo, Japan).

### Purification of recombinant TpPyShell1 protein

The expression plasmid for TpPyShell with an N-terminal His6-tag on pET28a was transformed into *Escherichia coli* strain BL21(DE3). Cells were cultured at 37°C in 6 L of LB medium containing 100 µg/mL Kanamycin. When the OD600 reached 0.5, an IPTG solution was added to a final concentration of 0.1 mM for induction of TpPyShell expression, and the culture was incubated overnight at 37°C. Cells were harvested by two rounds of centrifugation (6,000 rpm, 10 min, 4°C, JLA-9.1000 rotor; Beckman), resuspended in Buffer A containing 50 mM Tris-HCl pH8.0, 0.3 M NaCl, 1 mM EDTA and 1 mM DTT, and then frozen for storage at −80°C or used immediately for cell disruption. Frozen cells were resuspended in 200 mL of freshly prepared pre-cooled Buffer A with 0.25 mM PMSF and disrupted by sonication. After cell debris and undisrupted cells were removed by centrifugation (45,000 rpm, 30 min, 4°C, 70Ti rotor; Beckman), supernatant was applied to an open column with Ni-IMAC resin (BIO-RAD). The column was washed with wash buffer (50 mM Tris-HCl pH8.0, 0.3 M NaCl, 10 mM Imidazole, 1 mM EDTA, 1 mM DTT), and then TpPyShell protein was eluted from the column using elution buffer (50 mM Tris-HCl pH8.0, 0.3 M NaCl, 300 mM Imidazole, 1 mM EDTA and 1 mM DTT). TEV protease equivalent to 3% of the TpPyShell concentration was added to the fractionated solution. His-tagged TpPyShell together with TEV protease was dialyzed overnight at 4°C in SnakeSkin dialysis tube (Thermo Scientific) against dialysis buffer (50 mM Tris-HCl pH8.0, 0.3 M NaCl, 1 mM EDTA and 1 mM DTT). The dialyzed solution was applied to an open column with Ni-IMAC resin equilibrated by the dialysis buffer, and the His-tag free TpPyShell was eluted. Collected TpPyShell was applied to Superdex75 16/60 equilibrated with the dialysis buffer, and the fraction containing TpPyShell was collected. The fraction containing TpPyShell was further concentrated by centrifugation (4000 g, 15 min, 4°C, SX4400 rotor; Beckman) with Amicon Ultra (M.W.4000) to reach a TpPyShell concentration of 2.0 mg/mL.

### Cryo-EM grid prep and data acquisition

3.0 µL of TpPyShell solution was applied to a glow-discharged Quantifoil holey carbon grid (R1.2/1.3, Cu, 200 mech), blotted for 3.5 sec at 4°C and plunge-frozen into liquid ethane using a Vitrobot Mark IV (Thermo Fishrer Scientific). The grid was inserted into a Titan Krios (Thermo Fishrer Scientific) operating at an acceleration voltage of 300 kV and equipped with a Cs corrector (CEOS, GmbH). Images were recorded with a K3 direct electron detector (Gatan) in CDS mode with an energy filter at a slit width of 20 eV. Data were automatically collected using SerialEM software (Mastronarde, 2005) at a physical pixel size of 0.87 Å, with 52 frames at a dose of 0.96 e^-^/Å per frame, an exposure time of 2.63 sec per movie, and defocus ranging from −0.5 to −1.7 µm. A total of 5,951 movies were collected.

### Cryo-EM image processing and model building

The movie frames were subjected to beam-induced motion correction using MotionCorr2.1 (Zheng et al., 2017), and the contrast transfer function (CTF) was evaluated using Gctf (Zhang, 2016). Motion correction and CTF estimation were processed using RELION 3.1 (Zivanov et al., 2018). The motion corrected micrographs were imported into cryoSPARC ver.4.0.2 (Punjani et al., 2017), and approximately 500 particles were manually selected from 10 micrographs to perform two-dimensional (2D) classification. Using a good 2D class average image as a template, a total of 800,778 particle images were automatically picked from all micrographs in a filament tracer job and were extracted with a box size of 150 pixels with 4x binning. After two rounds of 2D classification, 299,145 particles were selected and extracted with a box size of 600 pixels. The re-extracted particles were subjected to refinement without helical parameters. Even after refinement, symmetry search for the helical parameters were not convincing. Therefore, preliminarily modelled structures were manually fitted into an EM density map to estimate two helical parameters, and the helical parameters were determined to be rise of 25 Å and twist of −30 degrees. Based on these helical parameters, 496,631 particle images were automatically picked from all micrographs and were extracted with a box size of 170 pixels with 4x binning using RELION 3.1. After 2D classification, selected particles were classified into 5 classes by 3D classification, as shown in Figure S5B. A total of 123,385 particles were re-extracted at a pixel size of 1.20 Å and subjected to five rounds of helical refinement, three rounds of CTF refinement, and Bayesian polishing. The final 3D refinement and post-processing yielded a map with global resolution of 3.0 Å, according Fourier shell correlation (FSC) with the 0.143 criterion (Figure S5E). The final refined values of helical rise and twist were 25.14 Å and −32.46 degrees, respectively. Local resolution was estimated using RELION 3.1 (Figure S5F). The processing workflow is outlined in Figure S5B.

The model of TpPyShell1 from Trp96 to Met316 (excluding the chloroplast targeting sequence) (Figures 3B, S5G) was built starting from the predicted AlphaFold2 model (Jumper et al., 2021). After manually fitting this predicted model into the EM density map using UCSF Chimera (Pettersen et al., 2004), each domain was manually remodeled and refined iteratively using COOT (Emsley et al., 2010), Phenix (Liebschner et al., 2019), and the Servalcat pipeline in REFMAC5 (Yamashita et al., 2021). All figures were prepared using UCSF ChimeraX (Goddard et al., 2018). The statics of the 3D reconstruction and model refinement are summarized in Table S5.

### Cryo-ET sample prep and data acquisition

*Thalasiossira pseudonana* and *Phaeodactylum tricornumtum* were grown in artificial sea water at 18°C and 40 μmol photons m^−2^ s^−1^ light without shaking. For the *Thalasiossira pseudonana m1* and *m2* mutants (ΔTpPyShell1/2), 5 mM NaCO_3_ was supplemented in the medium. Cells were sedimented at 800 × *g* for 5 min prior to vitrification, 4 µL of cell suspension was applied on 200-mesh R ¼ SiO_2_-film covered gold grids or 200-mesh R2/1 carbon-film covered copper grids (Quantifoil Micro Tools) (for *P. tricornutum* and *T. pseudonana* cells, respectively) and plunge frozen using a Vitrobot Mark IV (Thermo Fisher Scientific). EM grids were clipped into Autogrid supports (Thermo Fischer Scientific) and loaded into Aquilos 1 or 2 FIB-SEM instruments (Thermo Fischer Scientific), where they were thinned with a Gallium ion beam as previously described (Schaffer et al., 2017). The resulting EM grids with thin lamellae were transferred to a transmission electron microscope for tomographic imaging.

For *P. tricornutum*, cryo-ET data was acquired on a 300 kV Titan Krios G2 microscope (Thermo Fisher Scientific), equipped with a post-column energy filter (Quantum, Gatan) and a direct detector camera (K2 summit, Gatan) (“M1”), as well as on a 300 kV Titan Krios G3i microscope (Thermo Fisher Scientific) equipped with a BioQuantum post-column energy filter (Gatan) and a K3 direct electron detector (Gatan) (“M2”). For *T. pseudonana*, data was acquired on microscope M2, as well as on a 300 kV Titan Krios G4i microscope (Thermo Fisher Scientific), equipped with a post-column energy filter (Selectris X, Thermo Fisher Scientific) and a direct electron detector (Falcon 4, Thermo Fisher Scientific) (“M3”).

For microscopes M1 and M2, tilt-series were obtained using SerialEM 3.8 software (Mastronarde et al, 2005). In all cases, tilt-series were acquired using a dose-symmetric tilt scheme (Hagen et al, 2017), with 2° steps totaling 60 tilts per series. Each image was recorded in counting mode with ten frames per second. The target defocus of individual tilt-series ranged from −2 to −5 µm. Total dose per tilt series was approximately 120 e^-^/Å^2^. Image pixel sizes for microscopes M1 and M2 were 3.52 and 2.143 Å, respectively. For microscope M3, tilt-series were obtained using the Tomography 5.11 software (Thermo Fisher Scientific), using the same acquisition scheme as above, except for the use of multi-shot acquisition. Data was acquired in EER mode with a calibrated image pixel size of 2.93 Å.

### Cryo-ET data analysis

TOMOMAN Matlab scripts (https://github.com/williamnwan/TOMOMAN/; https://doi.org/10.5281/zenodo.4110737) (version 0.6.9) were used to preprocess the tomographic tilt series data. Raw frames were aligned using MotionCor2 (version 1.5.0) (Zheng et al., 2017), then tilt-series were dose-weighted (Grant and Grigorieff, 2015) followed by manual removal of bad tilts. The resulting tilt-series (binned 4 times, pixel sizes: 14.08 Å for M1, 8.57 Å for M2, 11.6 Å for M3) were aligned in IMOD (version 4.11) (Mastronarde and Held, 2017) using patch tracking and were reconstructed by weighted back projection. Cryo-CARE (version 0.2.1) (Buchholz et al., 2018) was applied on reconstructed tomogram pairs from odd and even raw frames to enhance contrast and remove noise. Snapshots of denoised tomograms were captured using the IMOD 3dmod viewer. Denoised tomograms were used as input for automatic segmentation using MemBrain-Seg (https://github.com/teamtomo/membrain-seg/). The resulting segmentations were manually curated in Amira (version 2021.2).

### Subtomogram averaging of *T. pseudonana* PyShell

For subtomogram averaging, only data from microscope M2 was used. Segmented surfaces corresponding to the PyShell were used as input to determine initial normal vectors in MemBrain’s point and normal sampling module (Lamm et al., 2022). Vectors were sampled densely on the surface with a spacing of 1.5 voxels at bin4 (8.572 Å/px). The resulting positions were used as initial coordinates to extract subvolumes (box size of 32 pixels) from bin4 tomograms corrected for the contrast transfer function (CTF) using phase flipping in IMOD. Starting from the normal vectors determined in MemBrain, multiple rounds of subtomogram alignment, averaging and classification were carried out in STOPGAP software (https://github.com/williamnwan/STOPGAP/; https://doi.org/10.5281/zenodo.3973664). False positives and poorly aligning particles were removed by classification steps. A second round of extraction, alignment and classification was performed at bin2 (4.286 Å/px), starting with the coordinates from the previous round of averaging in bin4. During particle alignment a maximum resolution of 16 Å was allowed to prevent overfitting.

To compare the *in vitro* TpPyShell1 SPA model and the *in situ* PyShell STA density map (Figure 3F), the SPA density map was unrolled from a tube to a flat sheet using the unroll command in ChimeraX (Goddard et al., 2018). SPA models of individual monomers were fitted in the unrolled density to form a flat model of the *in vitro* lattice. This model was then subsequently fitted in the *in situ* STA density map using rigid fitting in ChimeraX.

### Photosynthesis measurements

*T. pseudonana* WT, *m1*, and *m2* cells were cultured under LC and HC with 40 μmol photons m^−2^ s^−1^ light, harvested at the logarithmic growth phase, and resuspended in freshly prepared DIC-free F/2 artificial water. Chlorophyll *a* concentration of the samples was determined in 100% (v/v) methanol (Jeffrey and Haxo, 1968), and the cell samples were applied to an oxygen electrode (Hansatech, King’s Lynn, U.K.) at 10 µg chlorophyll *a* mL^−1^ in the DIC-free artificial sea water (pH 8.1). Simultaneous measurement of net O_2_ evolution rate with total DIC concentration in the sample mixture was achieved by an oxygen electrode and a gas-chromatography flame ionization detector (GC-8A, Shimadzu, Kyoto, Japan) during stepwise addition of NaHCO_3_, as previously reported (Kikutani et al., 2016). Measurements used constant actinic light of 900 µmol photons m^−2^ s^−1^. The photosynthetic parameters were calculated from the plot of O_2_ evolution rate against DIC concentration by curve fitting with the non-linear least squares method: *P*_max_, maximum net O_2_ evolution rate; *K*_0.5_, DIC concentration giving a half of *P*_max_; [DIC]_comp_, [DIC] giving no net O_2_ evolution; and APC, apparent photosynthetic conductance.

### FIB-SEM data acquisition and analysis

Sample preparation was performed as in (Uwizeye et al., 2021). FIB-SEM tomography was performed with a Zeiss CrossBeam 550 microscope, equipped with Fibics Atlas 3D software for tomography. The voxel size was 16×16×16 nm for the WT, 6×6×6 nm for the *m1* mutant, and 8×8×8 nm for the *m2* mutant. The whole volumes were imaged with an average of 300 frames for WT and 1000 frames for the mutants. Single cells were isolated by cropping in 3D using the open software Fiji (Schindelin et al., 2012). Image misalignment was corrected using the “StackReg” plugin in Fiji. We used 3D Slicer for segmentation and 3D reconstruction, and Meshlab to reduce noise and enhance contours of reconstructed objects. The quantitative measurements of chloroplasts and pyrenoids organelles (volume, diameter, sphericity) was implemented in python using libraries including trimesh, stl and scikit-image.

### Phylogenetic analysis

Homologs of *T. pseudonana* PyShell 1 protein (TpPyShell1) were retrieved from the National Centre for Biotechnology Information and the Marine Microbial Eukaryote Transcriptome Sequence Project (MMETSP) (Keeling et al., 2014). The highest scoring sequences per species were selected (E-value cutoff = 1e−35). Gaps and non-conserved regions were removed, and the protein sequences were subsequently aligned using Clustal Omega (Sievers et al., 2011). The alignment was used to generate a maximum likelihood tree, using IQTREE with standard settings and visualized with iTOL (Trifinopoulos et al., 2016; Letunic and Bork, 2021). Taxonomic distribution of the TpPyShell1 protein sequence in the ocean was queried against the Ocean Gene Atlas v2.0 webserver (https://tara-oceans.mio.osupytheas.fr) (Villar et al., 2018; Vernette et al., 2022).

## Data availability

Single particle cryo-EM maps (EMD-37751), cryo-ET subtomogram averages (EMD-18709), and cellular tomograms (EMD-18710 to EMD-18713) are available in the Electron Microscopy Data Bank. Atomic models of the PyShell structure are deposited at the Protein Data Bank (PDB-8WQP). Raw cryo-EM data (EMPIAR-11724) and raw cryo-ET data (EMPIAR-11747) used to generate the density maps are available on the Electron Microscopy Public Image Archive. The PyShell nucleotide sequences reported in this paper have been deposited in DDBJ/EMBL/GenBank under accession numbers XM_002179781, XM_002185069, OR682719, OR682720, XM_002292359, XM_002292147, XM_002292148. The proteomics data is available at ProteomeXchange (PXD041920) and jPOST (JPST001940). Strains and plasmids generated in this study are available from the corresponding authors upon request.

## Acknowledgements

We thank Vandana Tomar Deopa and Kohei Yoneda for helping establish the mutant strains, as well Nobuko Higashiuchi and Eri Nakayama for technical assistance. We thank Cécile Giustini for and Haythem Hsine for data analysis and strain maintenance, Glen Wheeler and Colin Brownlee (Marine Biological Association of the UK) for supporting the first cryo-ET investigations, and Caitlyn McCafferty for help with bioinformatic analysis. We thank Rikuya Morimoto, Kanae Fukushima and Taiki Fukuzawa for their help establishing the preparation of recombinant PyShell protein, as well as Atsunori Oshima and Yoshinori Fujiyoshi (Naogo University) for TEM instrumentation access at the beginning of the SPA cryo-EM study. We thank Guy Schoehn and Christine Moriscot (IBS/ISBG electron microscope facility; Grenoble Instruct-ERIC center) for access and assistance with FIB/SEM instrumentation. We thank Sagar Khavnekar, Jürgen Plitzko, and Wolfgang Baumeister (Max Plank Institute of Biochemistry), as well as Daniel Böhringer, Miroslav Peterek, and Bilal M. Qureshi (ETH ScopeM facility) for access to FIB and TEM instruments and assistance with cryo-ET data acquisition. Cryo-ET analysis was performed at the sciSCORE (http://scicore.unibas.ch/) scientific computing center at the University of Basel.

## Author contributions

Y.M. and B.E. initiated the project; Y.T. and N.M. identified PyShell proteins; H.N. and T.O. generated the PyShell mutants; H.M. performed qPCR analysis; G.S. and T.O. performed growth and photosynthesis analyses; N.M. and R.O. analyzed the localization of GFP-fused PyShell proteins; H.N. and M.D. performed bioinformatic analyses; A.T. performed immunoelectron microscopy; R.T. expressed and purified recombinant TpPyShell1 protein; C.G. confirmed the tube structure of TpPyShell1; A.K. prepared cryo-EM grids, acquired data and calculated the SPA density map; G.K. performed modeling and refinement; S.F. and W.W. acquired cryo-ET data; M.D, L.L., and R.R. analyzed cryo-ET data and performed subtomogram averaging; S.F. and B.G. performed sample preparation for FIB-SEM imaging; P.-H.J. acquired FIB-SEM tomograms; S.F. and C.U. performed FIB-SEM tomogram segmentation and analysis; G.S., M.D., S.F., G.K., G.F., B.E., and Y.M. wrote the manuscript with support from all other authors.

## Supplementary information

is available for this paper. Supplemental Tables S1-S5

Supplemental Figures S1-S6

Supplemental Movies 1-2 (to be submitted with revised manuscript)

Correspondence and requests for materials should be addressed to G.K., B.E., and Y.M.

## Abbreviations

CA: carbonic anhydrase
CCM: CO_2_-concentrating mechanism
cryo-ET: cryo-electron tomography
DIC: dissolved inorganic carbon
EPYC1: essential pyrenoid component 1
FBA: fructose 1,6-bisphosphate aldolase
FIB: focused ion beam
GC-FID: gas-chromatography flame ionization detector
HC: high (1%) CO_2_
LC: low (0.04%) CO_2_
LC-MS/MS: liquid chromatography coupled with tandem mass spectrometry
pAA: photo-reactive amino acids
PyShell: pyrenoid shell
Rubisco: ribulose 1,5-bisphosphate carboxylase/oxygenase
RbcL: Rubisco large subunit
STA: subtomogram averaging
SPA: single particle analysis.

---

**Figure S1.**
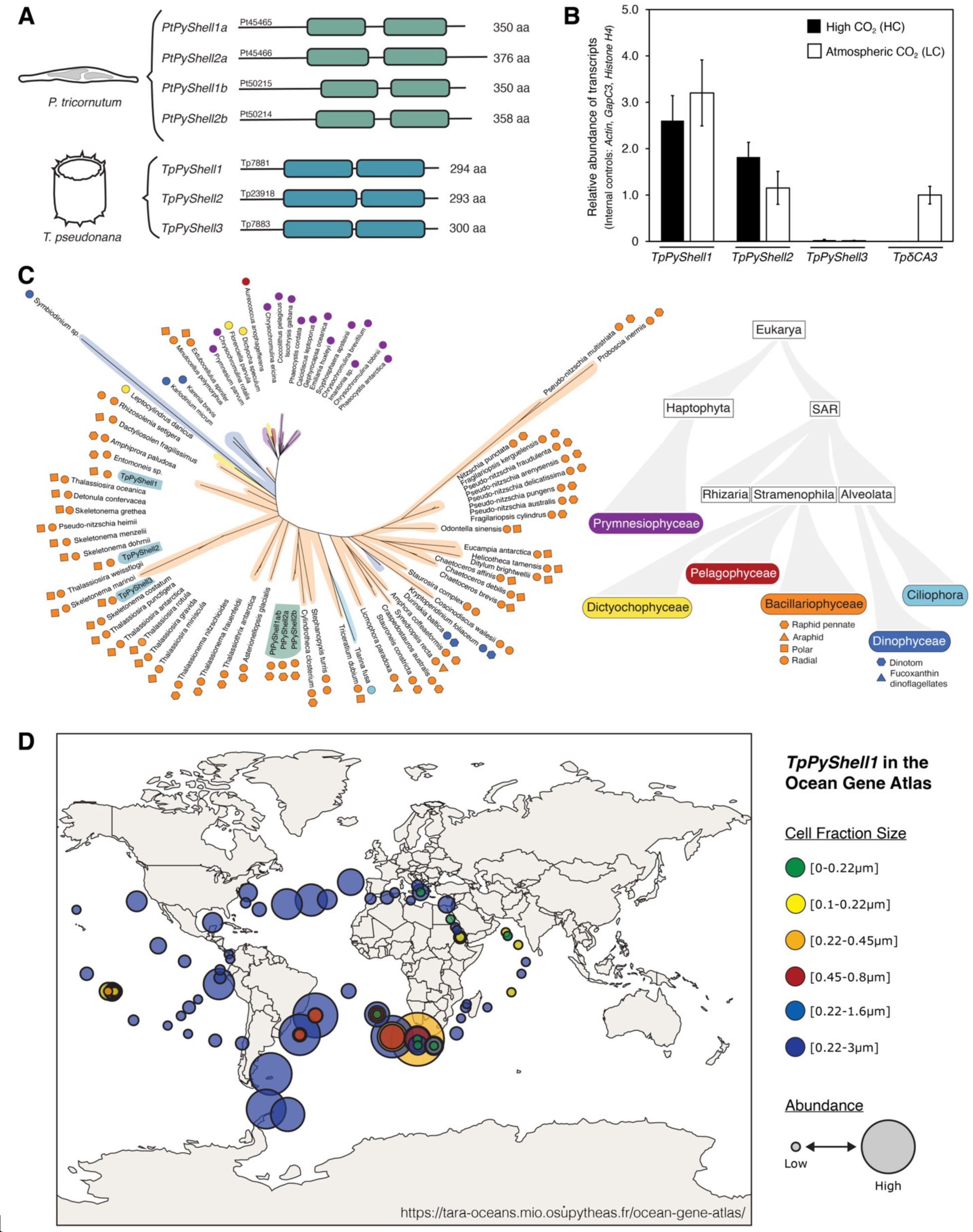
PyShell genes, expression, and phylogeny. **(A)** Domain architecture of the PyShell genes in *P. tricornutum* and *T. pseudonana*. The conserved CR1 and CR2 domains are indicated with colored bars. **(B)** Expression of the three PyShell genes identified in *T. pseudonana* under both normal atmosphere (LC, 0.04% CO_2_) and high CO_2_ (HC, 1%), compared to the expression of the CCM-induced carbonic anhydrase δCA3. **(C)** Maximum likelihood unrooted gene tree of TpPyshell1 (left), constructed with IQ-TREE, and an algal phylogenetic tree (right). The color in the phylogenetic tree indicates the clade to which each species in the gene tree belongs. Shapes and colors in the gene tree correspond to clades in the phylogenetic tree. The PyShell genes of *T. pseudonana* and *P. tricornutum* described in this study (TpPyshell1, 2, 3; PtPyShell1a/1b, 2a/2b) are highlighted in blue and teal, respectively. **(D)** Global distribution of TpPyShell1 homologous transcripts in fractions from *Tara* Oceans sampling (de Vargas et al., 2015), identified by searching the Ocean Gene Atlas v2.0 (Vernette et al., 2022).

**Figure S2.**
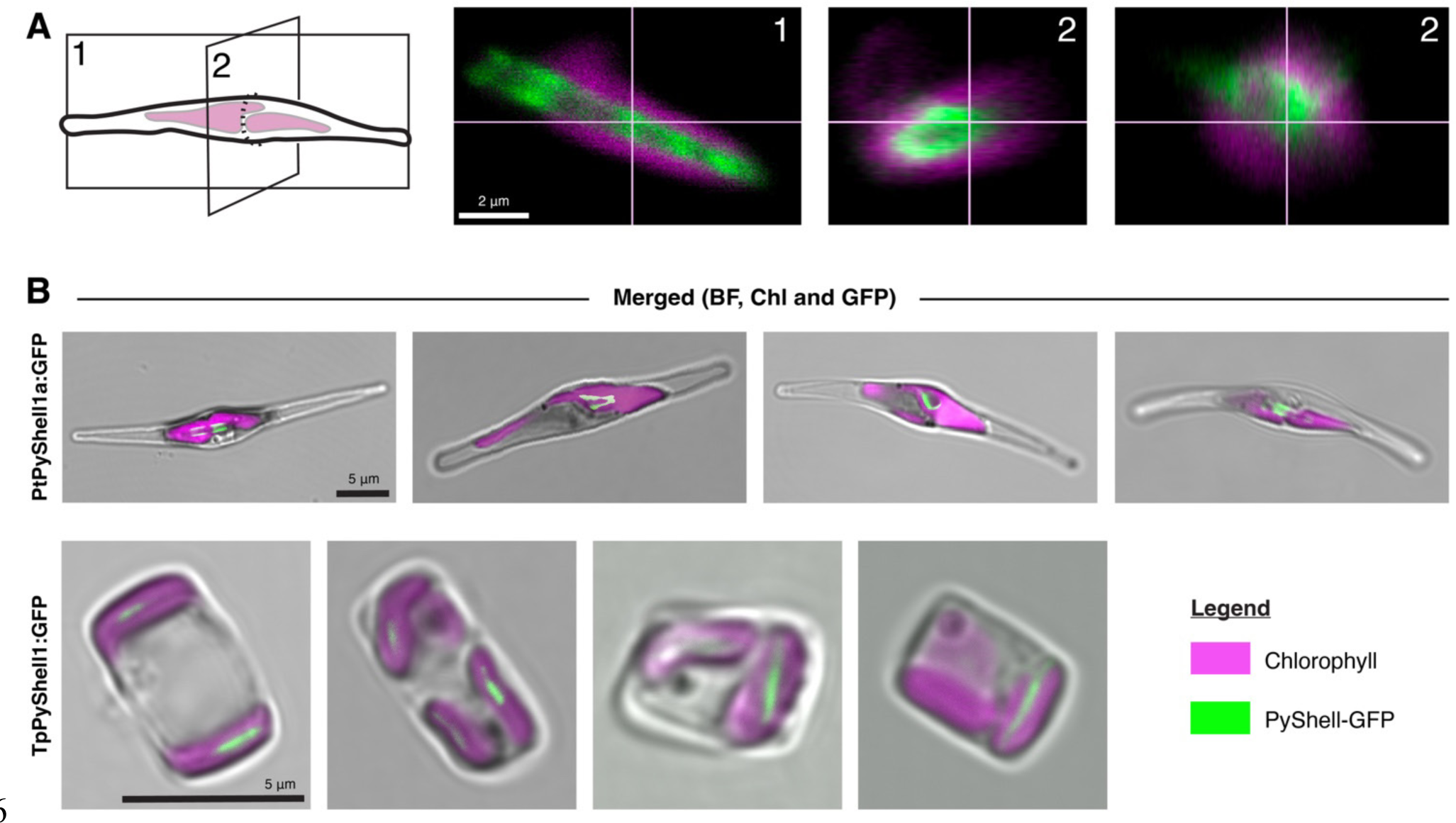
Additional fluorescence images of the PyShell in *P. tricornutum* and *T. pseudonana*. (**A)** 3D confocal images of PtPyShell1a:GFP in *P. tricornutum*. Diagram on the left shows the orientations of the virtual slices displayed on the right. **(B)** Maximum intensity projections of PtPyShell1a:GFP in *P. tricornutum* (top row) and TpPyShell1:GFP in *T. pseudonana* (bottom row). Grey: brightfield (BF), magenta: chlorophyl autofluorescence (Chl), green: GFP. Scale bars: 2 µm in A, 5 µm in B. Accompanies Fig. 1D.

**Figure S3.**
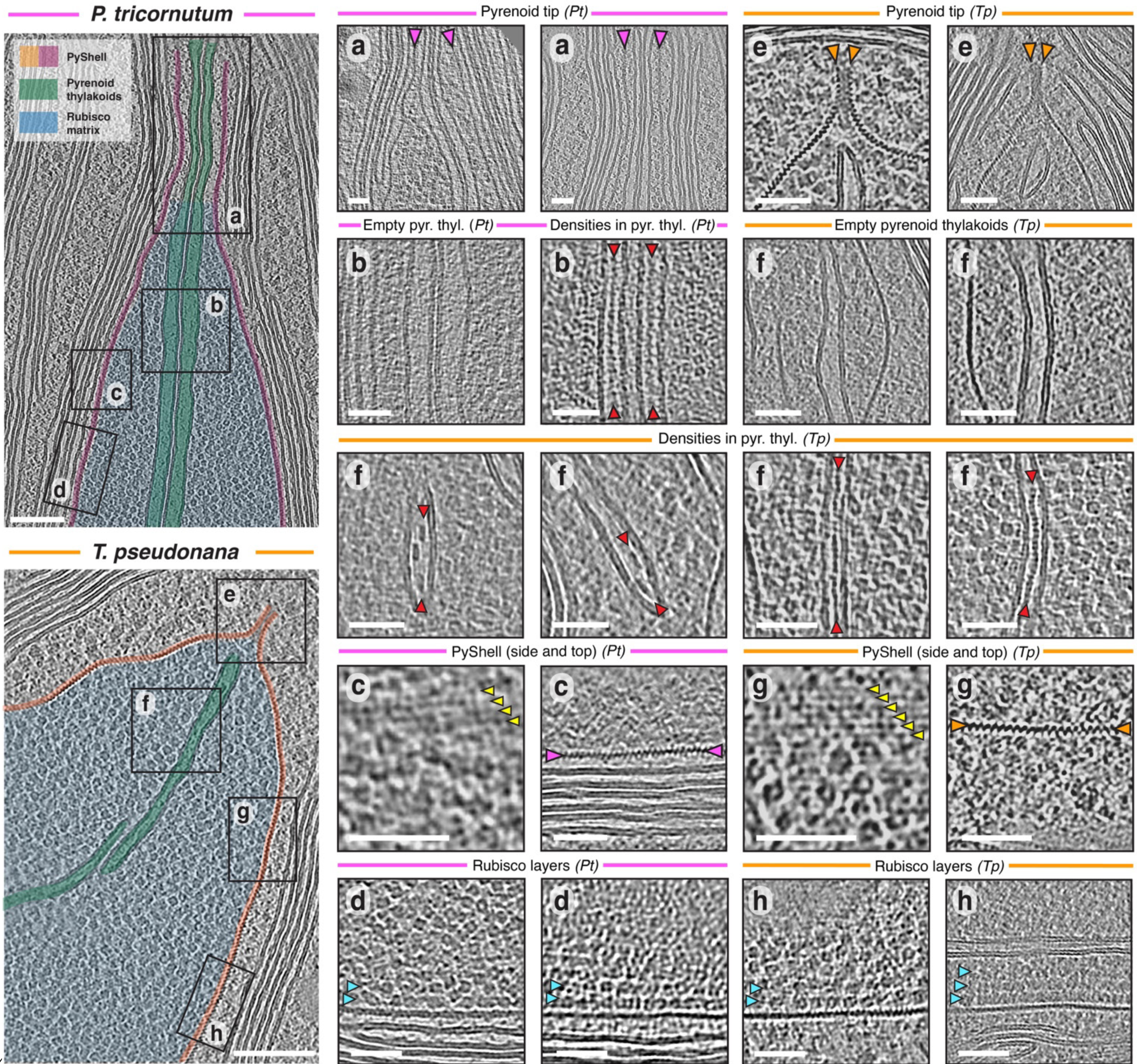
Additional cryo-ET images of pyrenoids inside native *P. tricornutum* and *T. pseudonana* cells. Left panels show pyrenoid overviews of *P. tricornutum* (pink) and *T. pseudonana* (orange), with lettered boxes indicating corresponding pyrenoid regions detailed in panels to the right. For *P. tricornutum* and *T. pseudonana*, respectively: **(A, E)** examples of the pyrenoid ends, which differ between species (PyShell: orange and pink arrowheads); **(B, F)** pyrenoid-traversing thylakoids, which sometimes have dense particles in the lumen (red arrowheads); **(C, G)** PyShell side (surface) and top (cross-section) views. Yellow arrowheads: major stripes of the PyShell lattice; **(D, H)** Ordered layers of Rubisco (blue arrowheads) adjacent to the PyShell. Scale bars: 100nm in overviews, 5 nm in all others. Accompanies Fig. 2A-I.

**Figure S4.**
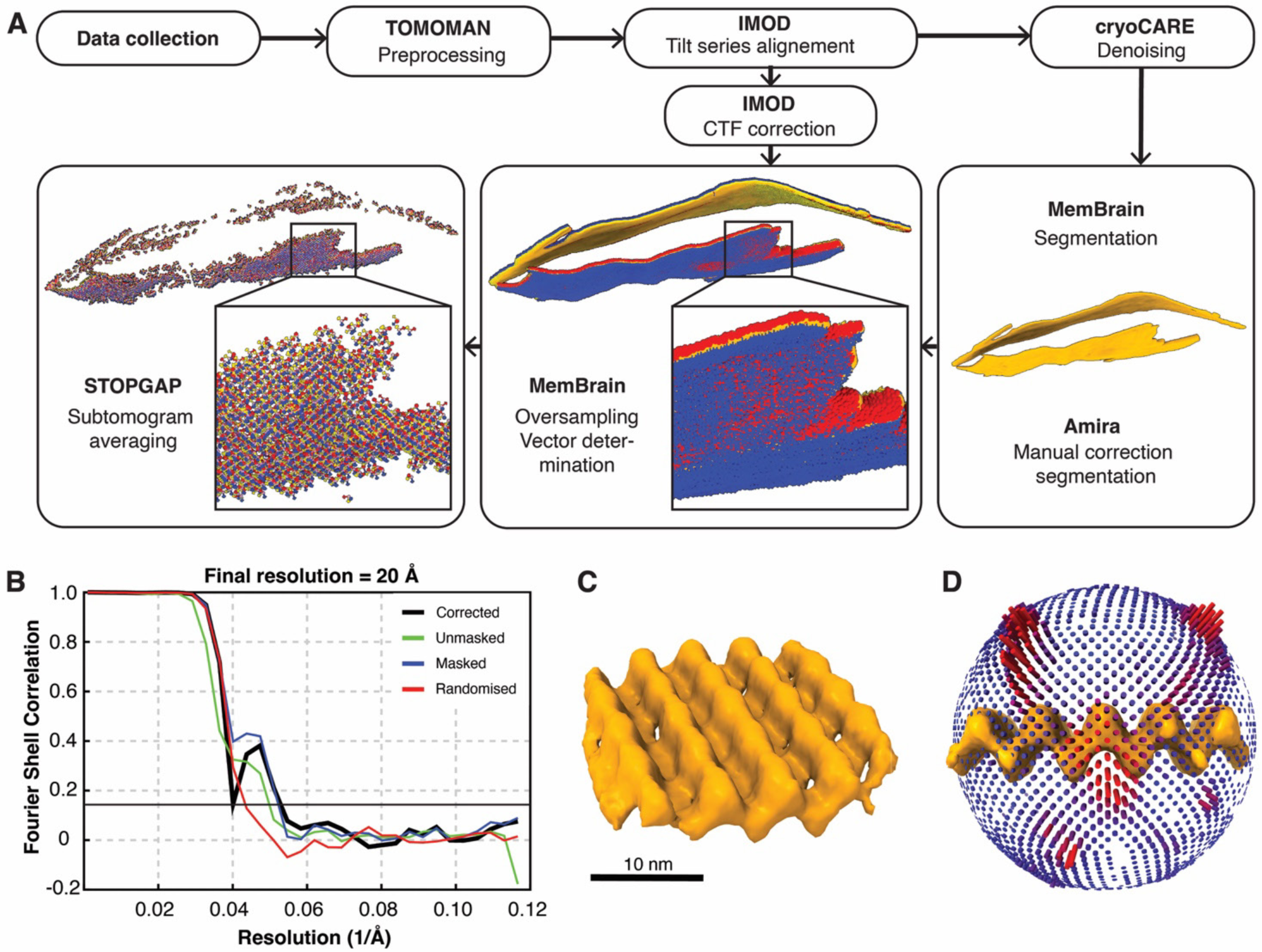
Cryo-ET subtomogram averaging supplement. **(A)** Cryo-ET data processing workflow. Determination of initial and final coordinates and vectors (blue/red/yellow arrows) is shown for an example tomogram. Coordinates were initially oversampled along the PyShell segmentation in MemBrain. After subtomogram averaging in STOPGAP, the coordinates converged to the repeat of the PyShell lattice. Only one subvolume per coordinate was retained in the final average. **(B)** FSC resolution determination of the resulting STA map, using the 0.143 cutoff. **(C)** Inclined view of the PyShell STA density map. Scale bar: 10 nm. **(D)** Angular distribution of particles contributing to the STA map. Red: more populated orientations, blue: less populated orientations.

**Figure S5.**
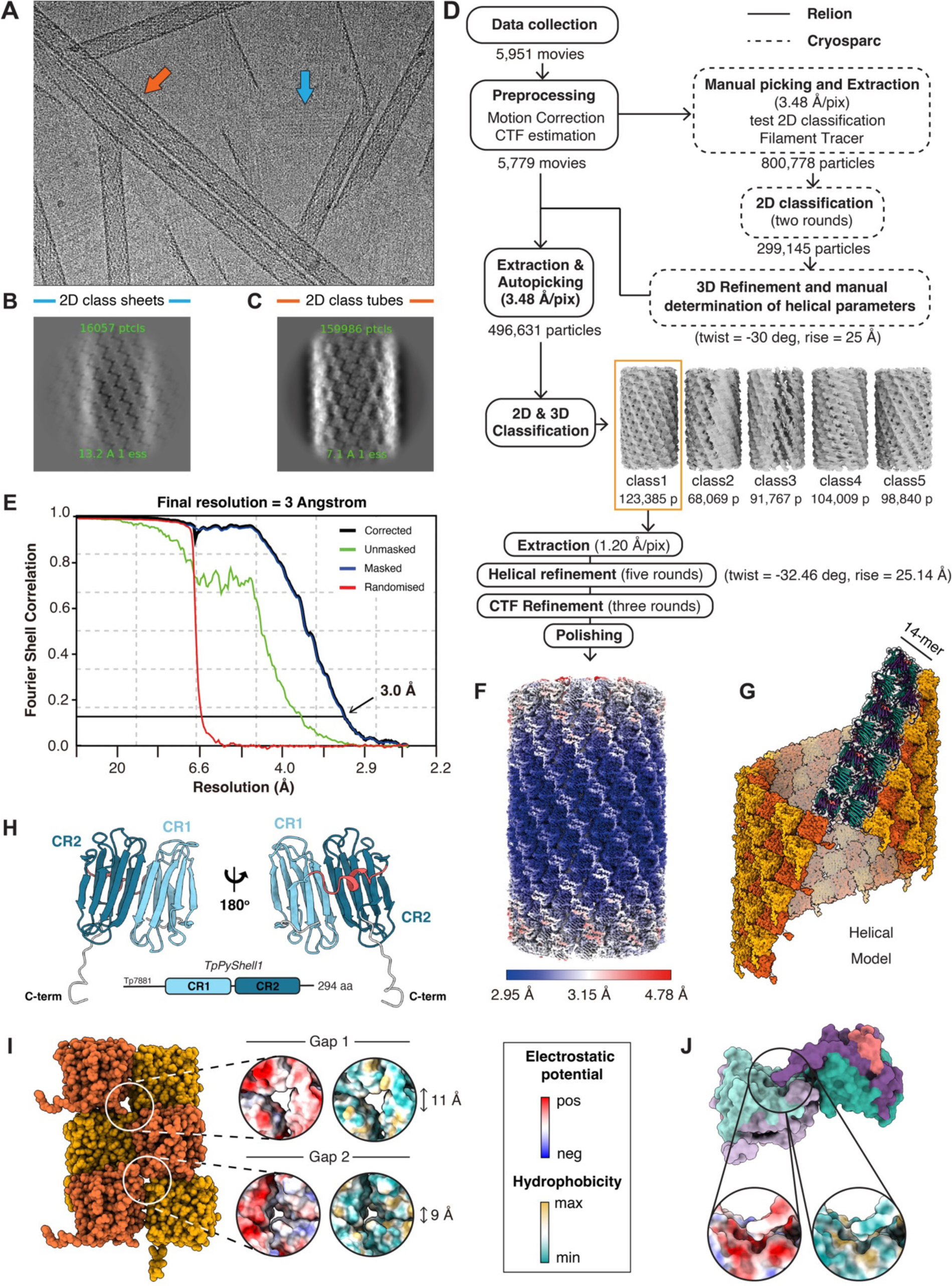
Single particle cryo-EM supplement. **(A)** Example micrograph. Sheets and tubes are indicated with blue and orange arrows, respectively. **(B)** 2D class average obtained for the TpPyShell1 sheets. **(C)** 2D class average obtained for the TpPyShell1 tubes. The homodimers of PyShell proteins can be seen in both the sheet and tube assemblies. **(D)** Cryo-EM processing workflow for high-resolution helical reconstruction of the TpPyShell1 tubes, displaying the five classes resulting from 3D classification. **(E)** FSC resolution determination of the resulting SPA map, using the 0.143 cutoff. **(F)** The EM density map displayed with local resolution. **(G)** The resulting helical protein model (orange and yellow surface representation) that was built using a 14mer repeating unit (purple and teal cartoon representation). **(H)** The internal pseudo-two-fold symmetry the TpPyShell1 monomer, subdividing the β-sheets between the β-5/-6 and β-13/-14 strands. Conserved regions CR1 and CR2 are colored (shades of blue), along with the short α-helix connecting CR1 with CR2 (pink). **(I)** Two potential gaps are present within the TpPyShell1 lattice. Orange and yellow surface representations correspond to the inward and outward facing monomers of the homodimer repeating unit. Circle insets: electrostatic potential and hydrophobicity are displayed on surface representations. **(J)** The C-terminal domain of each PyShell monomer reaches towards a potential pocket on the neighboring PyShell monomer. Above: colors on surface representations correspond to Fig. 3E. One monomer is displayed in lighter colors. Circle insets: Electrostatic potential and hydrophobicity are displayed on surface representations.

**Figure S6.**
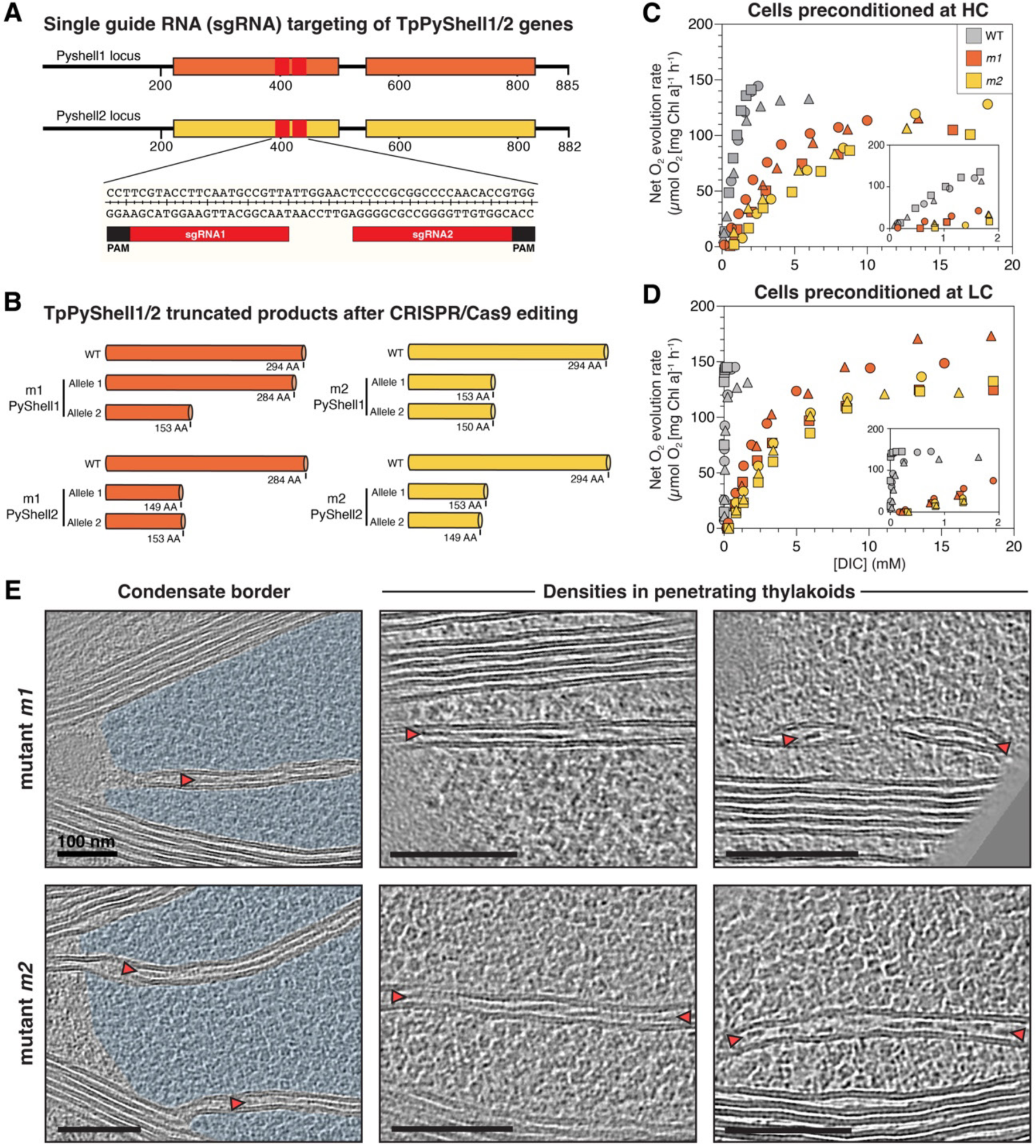
Generation of ΔTpPyPhell1/2 mutants, O_2_ evolution, and additional cryo-ET. **(A)** Schematic representation of simultaneous CRISPR/Cas9 targeting in both TpPyShell loci. **(B)** Resulting protein products in the mutant strains *m1* and *m2* at both loci. **(C, D)** Dependence of photosynthetic activity (measured by O_2_ evolution under 900 µmol photons m^−2^ s^−1^ constant actinic light) on DIC concentration (set by supplementing with bicarbonate) in WT cells (grey), *m1* (orange), and *m2* (yellow). Different symbols: three independent experiments. Cells were either preconditioned in HC or LC conditions; the WT preconditioned in LC has a more robust O_2_ evolution response in low DIC concentrations due to full activation of its CCM (compare inset panels). **(E)** Additional cryo-ET of Rubisco condensates in *m1* and *m2* cells. Light blue: Rubisco matrix; red arrowheads: densities in thylakoid lumen. Scale bars: 100 nm.

**Table S1.**
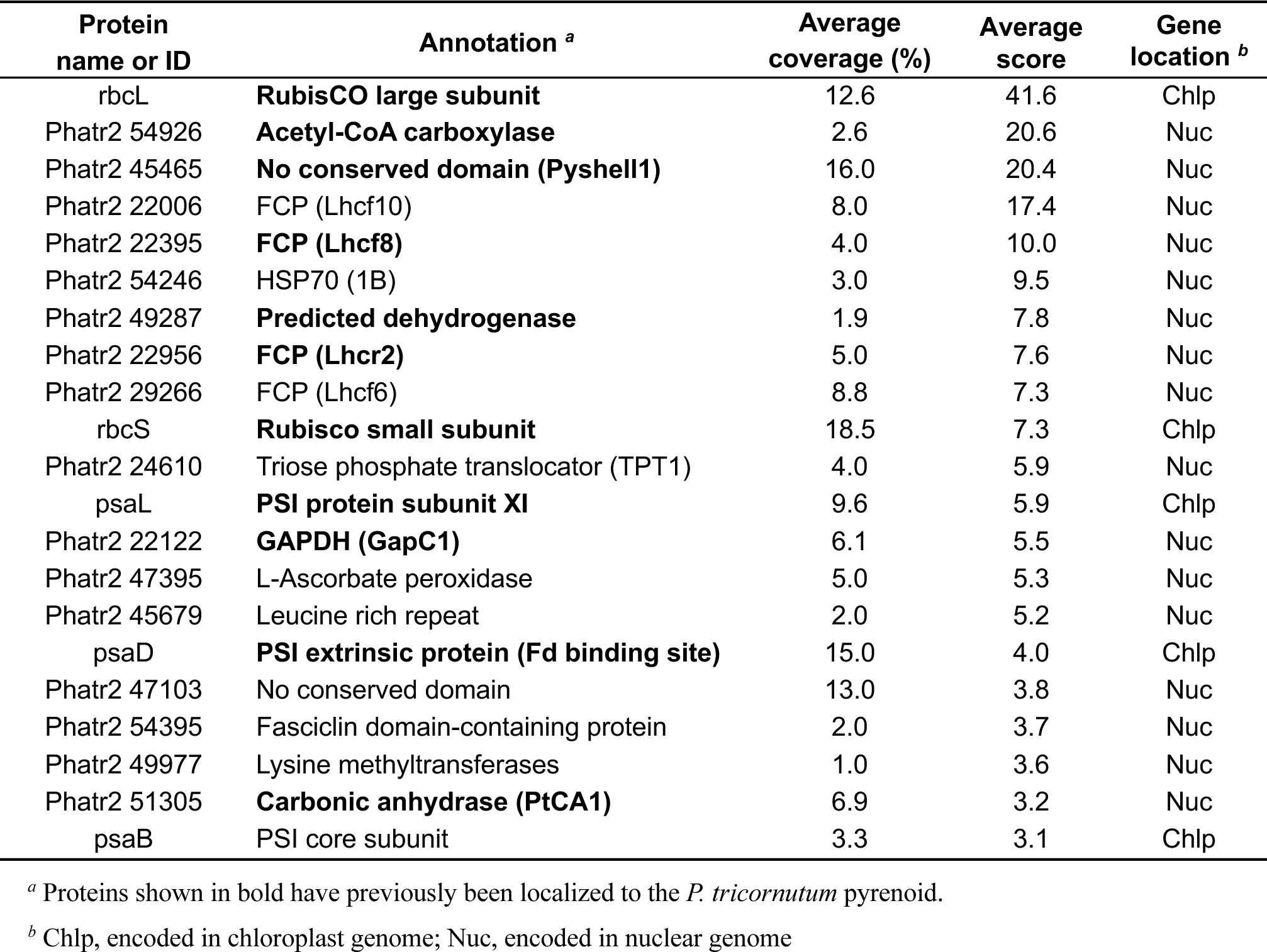
**List of candidate pyrenoid proteins in *P. tricornutum* identified by direct analysis of the stacking gel.** These “gel digestion” results correspond to “Procedure A” in Fig. 1A.

**Table S2.** List of candidate pyrenoid proteins in *P. tricornutum* identified in Rubisco-enriched fractions obtained by sucrose density gradient centrifugation. These “solution digestion” results correspond to “Procedure B” in Fig. 1A.

| Protein name or ID | Annotation <sup>a</sup> | Average coverage (%) | Average score | Gene location <sup>b</sup> |
| --- | --- | --- | --- | --- |
| rbcL | **RubisCO large subunit** | 16.8 | 71.8 | Chlp |
| Phatr2 51305 | **Carbonic anhydrase (PtCA1)** | 21.0 | 32.9 | Nuc |
| Phatr2 25168 | FCP (Lhcf4) | 21.0 | 22.8 | Nuc |
| psaF | PSI subunit (Cyt c<sub>6</sub> binding site) | 12.0 | 19.9 | Chlp |
| rbcS | **Rubisco small subunit** | 15.1 | 16.6 | Chlp |
| Phatr2 45465 | **No conserved domain (Pyshell1)** | 17.0 | 11.2 | Nuc |
| psbV | Cytochrome c<sub>550</sub> (PSII Mn-cluster) | 18.2 | 10.8 | Chlp |
| Phatr2 22122 | **GAPDH (GapC1)** | 7.0 | 9.8 | Nuc |
| Phatr2 13877 | FCP 47485 | 17.7 | 8.8 | Nuc |
| Phatr2 20331 | PsbO | 6.6 | 8.7 | Nuc |
| atpA | ATP synthase CF1 alpha subunit | 6.0 | 8.3 | Chlp |
| Phatr2 34919 | No conserved domain | 0.0 | 8.0 | Nuc |
| psbD | D2 protein | 5.6 | 7.7 | Chlp |
| Phatr2 14386 | FCP (Lhcr14) | 14.6 | 6.8 | Nuc |
| psaC | PSI extrinsic protein (Fd binding site) | 42.0 | 6.6 | Chlp |
| Phatr2 22956 | **FCP (Lhcr2)** | 8.0 | 6.4 | Nuc |
| Phatr2 26293 | PSII 12 kDa extrinsic protein (PsbU) | 10.0 | 6.3 | Nuc |
| atpB | FoF1 ATPase beta-subunit | 8.4 | 5.5 | Chlp |
| psaD | **PSI extrinsic protein (Fd binding site)** | 21.3 | 5.5 | Chlp |
| Phatr2 22395 | **FCP (Lhcf8)** | 11.8 | 5.4 | Nuc |
| Phatr2 51230 | FCP (Lhcf11) | 8.4 | 4.8 | Nuc |
| Phatr2 44488 | DNA polymerase III, delta subunit | 3.0 | 4.7 | Nuc |
| Phatr2 49287 | **Predicted dehydrogenase** | 1.5 | 4.7 | Nuc |
| Phatr2 25893 | FCP (Lhcf14) | 7.3 | 4.4 | Nuc |
| psbC | CP43 (Core antenna of PSII) | 3.2 | 4.1 | Chlp |
| Phatr2 30648 | FCP (Lhcf11) | 14.0 | 4.1 | Nuc |
| Phatr2 44056 | Cytochrome c<sub>553</sub> (PetJ) | 14.0 | 4.0 | Nuc |
| psaL | **Photosystem I protein subunit XI** | 13.6 | 4.0 | Chlp |
| psbA | D1 protein | 3.0 | 3.7 | Chlp |
| Phatr2 54926 | **Acetyl-CoA carboxylase** | 1.3 | 3.7 | Nuc |
| psbB | CP47 (Core antenna of PSII) | 9.0 | 3.6 | Chlp |
| Phatr2 20657 | FoF1 ATP synthase, gamma subunit | 4.0 | 3.6 | Nuc |
| Phatr2 46917 | No conserved domain | 6.0 | 3.5 | Nuc |
| psbE | Cytochrome b<sub>559</sub> | 23.7 | 3.5 | Chlp |
| ycf41 | Hypothetical chloroplast reading frame 41 | 13.0 | 3.3 | Chlp |
| Phatr2 42663 | No conserved domain | 1.3 | 3.2 | Nuc |

<sup>a</sup> Proteins shown in bold have previously been localized to the *P. tricornutum* pyrenoid.
<sup>b</sup> Chlp, encoded in chloroplast genome; Nuc, encoded in nuclear genome

**Table S3.** Photosynthetic parameters in *T. pseudonana* wild-type cells and PyShell mutants. *P*_max_, maximum net photosynthetic O_2_ evolution rate (µmol mg^−1^ chlorophyll *a* h^−1^); *K*_0.5_, [DIC] (mM) giving a half of *P*_max_; [DIC]_comp_, [DIC] (mM) giving no net O_2_ evolution; and APC, apparent photosynthetic conductance (µmol mg^−1^ chlorophyll *a* h^−1^ mM^−1^ [DIC]). Data are shown as the mean ± the standard deviation (*n* = 3, biological replicates).

| Pre-conditioning | Strains | $P_{\max}$ | $K_{0.5}$ | $[DIC]_{\text{comp}}$ | APC |
| --- | --- | --- | --- | --- | --- |
| Air-level CO<sub>2</sub> (LC) | WT | 142 ± 13 | 0.046 ± 0.019 | 0.014 ± 0.011 | 2800 ± 1000 |
|  | *m1* | 151 ± 26 | 2.4 ± 0.5 | 0.28 ± 0.08 | 50 ± 10 |
|  | *m2* | 137 ± 12 | 3.3 ± 0.6 | 0.35 ± 0.03 | 33 ± 5 |
| High CO<sub>2</sub> (HC) | WT | 157 ± 19 | 0.82 ± 0.06 | 0.07 ± 0.04 | 130 ± 20 |
|  | *m1* | 119 ± 3 | 3.0 ± 0.8 | 0.37 ± 0.16 | 36 ± 6 |
|  | *m2* | 123 ± 15 | 5.0 ± 0.8 | 0.58 ± 0.20 | 29 ± 7 |

**Table S4.**
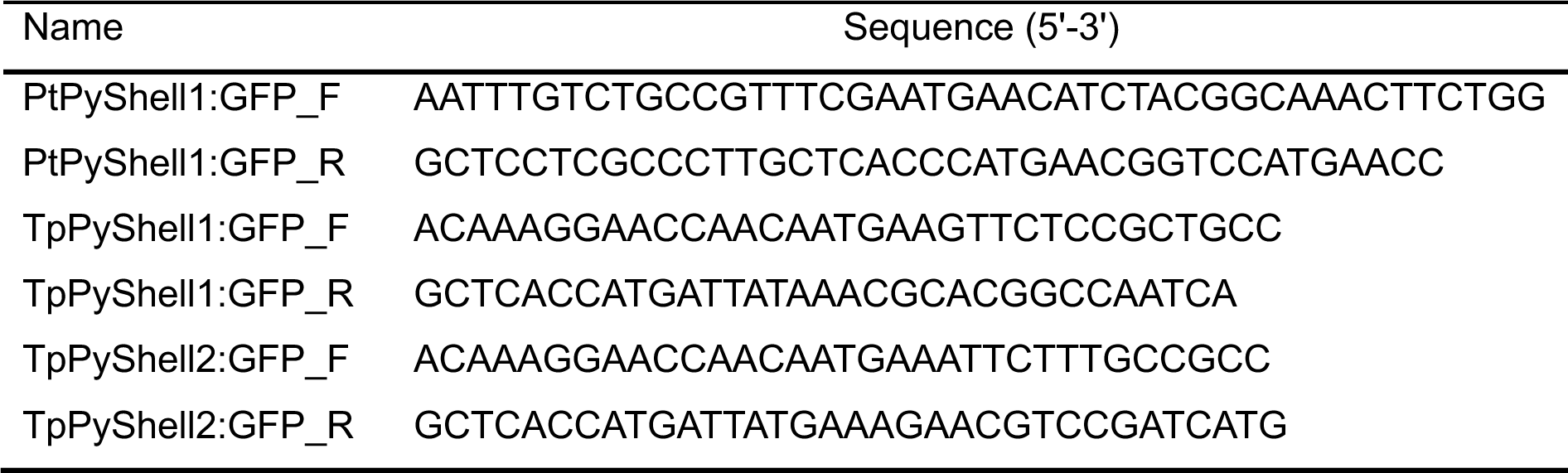
Primers used for expression of GFP fusion proteins in *P. tricornutum* and *T. pseudonana*. Corresponds to the section “Expression of GFP fusion proteins in *P. tricornutum* and *T. pseudonana*” in the Materials and Methods.

**Table S5.**
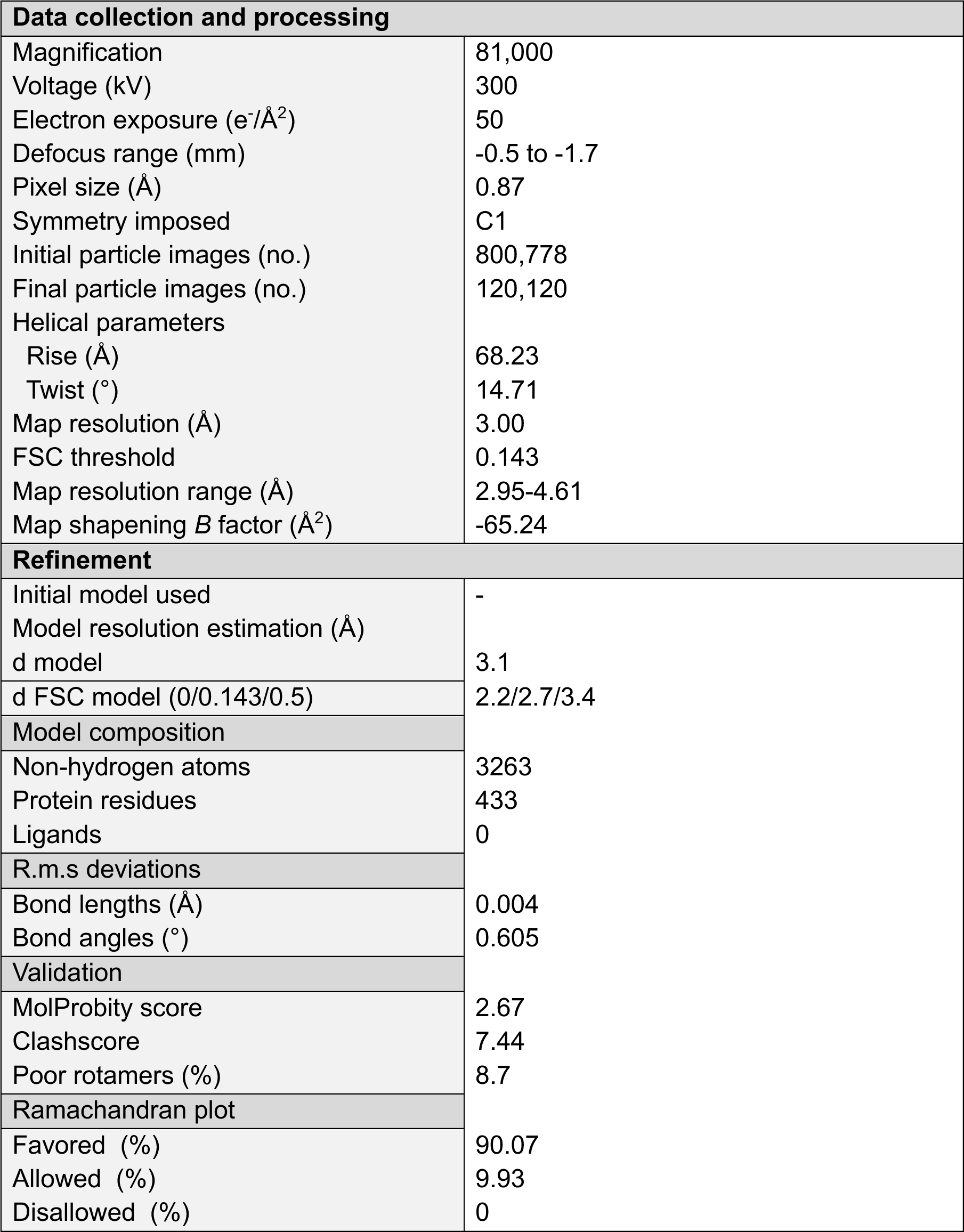
Cryo-EM data collection, refinement and validation statistics.

## References

1. Allen, A.E., Moustafa, A., Montsant, A., Eckert, A., Kroth, P.G., and Bowler, C. (2011). Evolution and functional diversification of fructose bisphosphate aldolase genes in photosynthetic marine diatoms. Molecular Biology and Evolution. 29: 367–379.

2. Atkinson, N., Mao, Y., Chan, K.X., and McCormick, A.J. (2020). Condensation of Rubisco into a proto-pyrenoid in higher plant chloroplasts. Nature Communications. 11: 6303.

3. Barrett, J., Girr, P., and Mackinder, L.C. (2021). Pyrenoids: CO_2_-fixing phase separated liquid organelles. Biochimica Et Biophysica Acta (BBA)-Molecular Cell Research. 1868: 118949.

4. Bedoshvili, Y.D., Popkova, T.P., and Likhoshway, Y.V. (2009). Chloroplast structure of diatoms of different classes. Cell and Tissue Biology. 3: 297–310.

5. Borden, J.S., and Savage, D.F. (2021). New discoveries expand possibilities for carboxysome engineering. Curr Opin Microbiol. 61: 58–66.

6. Borkhsenious, O.N., Mason, C.B., and Moroney, J.V. (1998). The intracellular localization of ribulose-1,5-bisphosphate Carboxylase/Oxygenase in chlamydomonas reinhardtii. Plant Physiol. 116: 1585–1591.

7. Buchholz, T.-O., Jordan, M., Pigino, G., and Jug, F. (2018). Cryo-CARE: Content-Aware Image Restoration for Cryo-Transmission Electron Microscopy Data. In arXiv e-prints, pp. arXiv:1810.05420.

8. Cai, F., Menon, B.B., Cannon, G.C., Curry, K.J., Shively, J.M., and Heinhorst, S. (2009). The pentameric vertex proteins are necessary for the icosahedral carboxysome shell to function as a CO_2_ leakage barrier. PLoS One. 4: e7521.

9. Cai, F., Sutter, M., Cameron, J.C., Stanley, D.N., Kinney, J.N., and Kerfeld, C.A. (2013). The structure of CcmP, a tandem bacterial microcompartment domain protein from the β-carboxysome, forms a subcompartment within a microcompartment. J Biol Chem. 288: 16055–16063.

10. Cole, J.J., Hararuk, O., and Solomon, C.T. (2021). Chapter 7 - The Carbon Cycle: With a Brief Introduction to Global Biogeochemistry. In Fundamentals of Ecosystem Science (Second Edition), K.C. Weathers, D.L. Strayer, and G.E. Likens, eds. (Academic Press), pp. 131–160.

11. de Vargas, C., Audic, S., Henry, N., Decelle, J., Mahé, F., Logares, R., Lara, E., Berney, C., Le Bescot, N., Probert, I., Carmichael, M., Poulain, J., Romac, S., Colin, S., Aury, J.-M., Bittner, L., Chaffron, S., Dunthorn, M., Engelen, S., Flegontova, O., Guidi, L., Horák, A., Jaillon, O., Lima-Mendez, G., Lukeš, J., Malviya, S., Morard, R., Mulot, M., Scalco, E., Siano, R., Vincent, F., Zingone, A., Dimier, C., Picheral, M., Searson, S., Kandels-Lewis, S., Acinas, S.G., Bork, P., Bowler, C., Gorsky, G., Grimsley, N., Hingamp, P., Iudicone, D., Not, F., Ogata, H., Pesant, S., Raes, J., Sieracki, M.E., Speich, S., Stemmann, L., Sunagawa, S., Weissenbach, J., Wincker, P., and Karsenti, E. (2015). Eukaryotic plankton diversity in the sunlit ocean. Science. 348: 1261605.

12. Dou, Z., Heinhorst, S., Williams, E.B., Murin, C.D., Shively, J.M., and Cannon, G.C. (2008). CO_2_ fixation kinetics of Halothiobacillus neapolitanus mutant carboxysomes lacking carbonic anhydrase suggest the shell acts as a diffusional barrier for CO_2_. J Biol Chem. 283: 10377–10384.

13. Emsley, P., Lohkamp, B., Scott, W.G., and Cowtan, K. (2010). Features and development of Coot. Acta Crystallogr D Biol Crystallogr. 66: 486–501.

14. Engel, B.D., Schaffer, M., Kuhn Cuellar, L., Villa, E., Plitzko, J.M., and Baumeister, W. (2015). Native architecture of the Chlamydomonas chloroplast revealed by in situ cryo-electron tomography. Elife. 4: e04889.

15. Falkowski, P.G., Barber, R.T., and Smetacek, V. (1998). Biogeochemical controls and feedbacks on ocean primary production. Science. 281: 200–206.

16. Faulkner, M., Szabó, I., Weetman, S.L., Sicard, F., Huber, R.G., Bond, P.J., Rosta, E., and Liu, L.N. (2020). Molecular simulations unravel the molecular principles that mediate selective permeability of carboxysome shell protein. Sci Rep. 10: 17501.

17. Fei, C., Wilson, A.T., Mangan, N.M., Wingreen, N.S., and Jonikas, M.C. (2022). Modelling the pyrenoid-based CO(2)-concentrating mechanism provides insights into its operating principles and a roadmap for its engineering into crops. Nat Plants. 8: 583–595.

18. Flori, S., Jouneau, P.-H., Bailleul, B., Gallet, B., Estrozi, L.F., Moriscot, C., Bastien, O., Eicke, S., Schober, A., Bártulos, C.R., Maréchal, E., Kroth, P.G., Petroutsos, D., Zeeman, S., Breyton, C., Schoehn, G., Falconet, D., and Finazzi, G. (2017). Plastid thylakoid architecture optimizes photosynthesis in diatoms. Nature Communications. 8: 15885.

19. Freeman Rosenzweig, E.S., Xu, B., Kuhn Cuellar, L., Martinez-Sanchez, A., Schaffer, M., Strauss, M., Cartwright, H.N., Ronceray, P., Plitzko, J.M., Forster, F., Wingreen, N.S., Engel, B.D., Mackinder, L.C.M., and Jonikas, M.C. (2017). The eukaryotic CO_2_-concentrating organelle is liquid-like and exhibits dynamic reorganization. Cell. 171: 148–162.e119.

20. Funke, R.P., Kovar, J.L., and Weeks, D.P. (1997). Intracellular carbonic anhydrase is essential to photosynthesis in Chlamydomonas reinhardtii at atmospheric levels of CO_2_. Demonstration via genomic complementation of the high-CO_2_-requiring mutant ca-1. Plant Physiol. 114: 237–244.

21. Giordano, M., Beardall, J., and Raven, J.A. (2005). CO_2_ concentrating mechanisms in algae: mechanisms, environmental modulation, and evolution. Annu Rev Plant Biol. 56: 99–131.

22. Goddard, T.D., Huang, C.C., Meng, E.C., Pettersen, E.F., Couch, G.S., Morris, J.H., and Ferrin, T.E. (2018). UCSF ChimeraX: Meeting modern challenges in visualization and analysis. Protein science: a publication of the Protein Society. 27: 14–25.

23. Grant, T., and Grigorieff, N. (2015). Measuring the optimal exposure for single particle cryo-EM using a 2.6 Å reconstruction of rotavirus VP6. eLife. 4: e06980.

24. Griffiths, D.J. (1970). The Pyrenoid. The Botanical Review. 36: 29–58.

25. Gruber, A., Vugrinec, S., Hempel, F., Gould, S.B., Maier, U.-G., and Kroth, P.G. (2007). Protein targeting into complex diatom plastids: functional characterisation of a specific targeting motif. Plant Molecular Biology. 64: 519–530.

26. Guillard, R.R.L. (1975). Culture of phytoplankton for feeding marine invertebrates. In Culture of marine invertebrate animals, W.L. Smith, and M.H. Chanley, eds. (Boston, MA: Springer US), pp. 29–60.

27. Guillard, R.R.L., and Ryther, J.H. (1962). Studies of marine planktonic diatoms: I. *Cyclotella nana* hustedt, and *Detonula confervacea* (cleve) gran. Canadian Journal of Microbiology. 8: 229–239.

28. Hanson, D.T., Franklin, L.A., Samuelsson, G., and Badger, M.R. (2003). The Chlamydomonas reinhardtii cia3 mutant lacking a thylakoid lumen-localized carbonic anhydrase is limited by CO_2_ supply to rubisco and not photosystem II function in vivo. Plant Physiol. 132: 2267–2275.

29. He, S., Chou, H.-T., Matthies, D., Wunder, T., Meyer, M.T., Atkinson, N., Martinez-Sanchez, A., Jeffrey, P.D., Port, S.A., Patena, W., He, G., Chen, V.K., Hughson, F.M., McCormick, A.J., Mueller-Cajar, O., Engel, B.D., Yu, Z., and Jonikas, M.C. (2020). The structural basis of Rubisco phase separation in the pyrenoid. Nature Plants. 6: 1480–1490.

30. Hennacy, J.H., and Jonikas, M.C. (2020). Prospects for Engineering Biophysical CO(2) Concentrating Mechanisms into Land Plants to Enhance Yields. Annu Rev Plant Biol. 71: 461–485.

31. Huang, J., Jiang, Q., Yang, M., Dykes, G.F., Weetman, S.L., Xin, W., He, H.L., and Liu, L.N. (2022). Probing the Internal pH and Permeability of a Carboxysome Shell. Biomacromolecules. 23: 4339–4348.

32. Hylton, R.K., and Swulius, M.T. (2021). Challenges and triumphs in cryo-electron tomography. iScience. 24: 102959.

33. Jeffrey, S., and Haxo, F. (1968). Photosynthetic pigments of symbiotic dinoflagellates (zooxanthellae) from corals and clams. The Biological Bulletin. 135: 149–165.

34. Jenks, A., and Gibbs, S.P. (2000). Immunolocalization and distribution of form II Rubisco in the pyrenoid and chloroplast stroma of *Amphidinium carterae* and form I Rubisco in the symbiont-derived plastids of *peridinium foliaceum* (Dinophyceae). Journal of Phycology. 36: 127–138.

35. Jumper, J., Evans, R., Pritzel, A., Green, T., Figurnov, M., Ronneberger, O., Tunyasuvunakool, K., Bates, R., Žídek, A., Potapenko, A., Bridgland, A., Meyer, C., Kohl, S.A.A., Ballard, A.J., Cowie, A., Romera-Paredes, B., Nikolov, S., Jain, R., Adler, J., Back, T., Petersen, S., Reiman, D., Clancy, E., Zielinski, M., Steinegger, M., Pacholska, M., Berghammer, T., Bodenstein, S., Silver, D., Vinyals, O., Senior, A.W., Kavukcuoglu, K., Kohli, P., and Hassabis, D. (2021). Highly accurate protein structure prediction with AlphaFold. Nature. 596: 583–589.

36. Karlsson, J., Clarke, A.K., Chen, Z.Y., Hugghins, S.Y., Park, Y.I., Husic, H.D., Moroney, J.V., and Samuelsson, G. (1998). A novel alpha-type carbonic anhydrase associated with the thylakoid membrane in Chlamydomonas reinhardtii is required for growth at ambient CO_2_. Embo j. 17: 1208–1216.

37. Keeling, P.J., Burki, F., Wilcox, H.M., Allam, B., Allen, E.E., Amaral-Zettler, L.A., Armbrust, E.V., Archibald, J.M., Bharti, A.K., Bell, C.J., Beszteri, B., Bidle, K.D., Cameron, C.T., Campbell, L., Caron, D.A., Cattolico, R.A., Collier, J.L., Coyne, K., Davy, S.K., Deschamps, P., Dyhrman, S.T., Edvardsen, B., Gates, R.D., Gobler, C.J., Greenwood, S.J., Guida, S.M., Jacobi, J.L., Jakobsen, K.S., James, E.R., Jenkins, B., John, U., Johnson, M.D., Juhl, A.R., Kamp, A., Katz, L.A., Kiene, R., Kudryavtsev, A., Leander, B.S., Lin, S., Lovejoy, C., Lynn, D., Marchetti, A., McManus, G., Nedelcu, A.M., Menden-Deuer, S., Miceli, C., Mock, T., Montresor, M., Moran, M.A., Murray, S., Nadathur, G., Nagai, S., Ngam, P.B., Palenik, B., Pawlowski, J., Petroni, G., Piganeau, G., Posewitz, M.C., Rengefors, K., Romano, G., Rumpho, M.E., Rynearson, T., Schilling, K.B., Schroeder, D.C., Simpson, A.G., Slamovits, C.H., Smith, D.R., Smith, G.J., Smith, S.R., Sosik, H.M., Stief, P., Theriot, E., Twary, S.N., Umale, P.E., Vaulot, D., Wawrik, B., Wheeler, G.L., Wilson, W.H., Xu, Y., Zingone, A., and Worden, A.Z. (2014). The Marine Microbial Eukaryote Transcriptome Sequencing Project (MMETSP): illuminating the functional diversity of eukaryotic life in the oceans through transcriptome sequencing. PLoS Biol. 12: e1001889.

38. Kikutani, S., Nakajima, K., Nagasato, C., Tsuji, Y., Miyatake, A., and Matsuda, Y. (2016). Thylakoid luminal θ-carbonic anhydrase critical for growth and photosynthesis in the marine diatom Phaeodactylum tricornutum. Proceedings of the National Academy of Sciences. 113: 9828–9833.

39. Klein, M.G., Zwart, P., Bagby, S.C., Cai, F., Chisholm, S.W., Heinhorst, S., Cannon, G.C., and Kerfeld, C.A. (2009). Identification and structural analysis of a novel carboxysome shell protein with implications for metabolite transport. J Mol Biol. 392: 319–333.

40. Kroth, P.G., and Matsuda, Y. (2022). Carbohydrate metabolism. In The Molecular Life of Diatoms (Springer), pp. 465–492.

41. Lacoste-Royal, G., and Gibbs, S.P. (1987). Immunocytochemical Localization of Ribulose-1,5-Bisphosphate Carboxylase in the Pyrenoid and Thylakoid Region of the Chloroplast of Chlamydomonas reinhardtii. Plant Physiol. 83: 602–606.

42. Lamm, L., Righetto, R.D., Wietrzynski, W., Pöge, M., Martinez-Sanchez, A., Peng, T., and Engel, B.D. (2022). MemBrain: A deep learning-aided pipeline for detection of membrane proteins in Cryo-electron tomograms. Comput Methods Programs Biomed. 224: 106990.

43. Larsson, A.M., Hasse, D., Valegård, K., and Andersson, I. (2017). Crystal structures of β-carboxysome shell protein CcmP: ligand binding correlates with the closed or open central pore. J Exp Bot. 68: 3857–3867.

44. Letunic, I., and Bork, P. (2021). Interactive Tree Of Life (iTOL) v5: an online tool for phylogenetic tree display and annotation. Nucleic Acids Research. 49: W293–W296.

45. Liebschner, D., Afonine, P.V., Baker, M.L., Bunkóczi, G., Chen, V.B., Croll, T.I., Hintze, B., Hung, L.W., Jain, S., McCoy, A.J., Moriarty, N.W., Oeffner, R.D., Poon, B.K., Prisant, M.G., Read, R.J., Richardson, J.S., Richardson, D.C., Sammito, M.D., Sobolev, O.V., Stockwell, D.H., Terwilliger, T.C., Urzhumtsev, A.G., Videau, L.L., Williams, C.J., and Adams, P.D. (2019). Macromolecular structure determination using X-rays, neutrons and electrons: recent developments in Phenix. Acta Crystallogr D Struct Biol. 75: 861–877.

46. Livak, K.J., and Schmittgen, T.D. (2001). Analysis of relative gene expression data using real-time quantitative PCR and the 2(-Delta Delta C(T)) Method. Methods. 25: 402–408.

47. Mackinder, L.C., Meyer, M.T., Mettler-Altmann, T., Chen, V.K., Mitchell, M.C., Caspari, O., Freeman Rosenzweig, E.S., Pallesen, L., Reeves, G., Itakura, A., Roth, R., Sommer, F., Geimer, S., Mühlhaus, T., Schroda, M., Goodenough, U., Stitt, M., Griffiths, H., and Jonikas, M.C. (2016). A repeat protein links Rubisco to form the eukaryotic carbon-concentrating organelle. Proc Natl Acad Sci U S A. 113: 5958–5963.

48. Mahinthichaichan, P., Morris, D.M., Wang, Y., Jensen, G.J., and Tajkhorshid, E. (2018). Selective Permeability of Carboxysome Shell Pores to Anionic Molecules. J Phys Chem B. 122: 9110–9118.

49. Mastronarde, D.N. (2005). Automated electron microscope tomography using robust prediction of specimen movements. J Struct Biol. 152: 36–51.

50. Mastronarde, D.N., and Held, S.R. (2017). Automated tilt series alignment and tomographic reconstruction in IMOD. Journal of Structural Biology. 197: 102–113.

51. Matsui, H., Hopkinson, B.M., Nakajima, K., and Matsuda, Y. (2018). Plasma Membrane-Type Aquaporins from Marine Diatoms Function as CO(2)/NH(3) Channels and Provide Photoprotection. Plant Physiol. 178: 345–357.

52. McGrath, J.M., and Long, S.P. (2014). Can the cyanobacterial carbon-concentrating mechanism increase photosynthesis in crop species? A theoretical analysis. Plant Physiol. 164: 2247–2261.

53. Melnicki, M.R., Sutter, M., and Kerfeld, C.A. (2021). Evolutionary relationships among shell proteins of carboxysomes and metabolosomes. Curr Opin Microbiol. 63: 1–9.

54. Meyer, M.T., Whittaker, C., and Griffiths, H. (2017). The algal pyrenoid: key unanswered questions. Journal of Experimental Botany. 68: 3739–3749.

55. Mocaer, K., Mizzon, G., Gunkel, M., Halavatyi, A., Steyer, A., Oorschot, V., Schorb, M., Le Kieffre, C., Yee, D.P., Chevalier, F., Gallet, B., Decelle, J., Schwab, Y., and Ronchi, P. (2023). Targeted volume correlative light and electron microscopy of an environmental marine microorganism. J Cell Sci. 136.

56. Morita, E., Kuroiwa, H., Kuroiwa, T., and Nozaki, H. (1997). High localization of ribulose-1,5-bisphosphate carboxulase/oxygenase in the pyrenoids of *Chlamydomonas reinhardtii* (Chlorophyta), as revealed by cryofixation and immunogold electron microscopy. Journal of phycology. 33: 68–72.

57. Motohashi, K. (2015). A simple and efficient seamless DNA cloning method using SLiCE from *Escherichia coli* laboratory strains and its application to SLiP site-directed mutagenesis. BMC Biotechnology. 15: 47.

58. Nawaly, H., Tanaka, A., Toyoshima, Y., Tsuji, Y., and Matsuda, Y. (2023). Localization and characterization θ carbonic anhydrases in Thalassiosira pseudonana. Photosynth Res. 156: 217–229.

59. Nawaly, H., Tsuji, Y., and Matsuda, Y. (2020). Rapid and precise genome editing in a marine diatom, *Thalassiosira pseudonana* by Cas9 nickase (D10A). Algal Research. 47: 101855.

60. Nonoyama, T., Kazamia, E., Nawaly, H., Gao, X., Tsuji, Y., Matsuda, Y., Bowler, C., Tanaka, T., and G. Dorrell, R. (2019). Metabolic innovations underpinning the origin and diversification of the diatom chloroplast. Biomolecules. 9: 322.

61. Oh, Z.G., Ang, W.S.L., Poh, C.W., Lai, S.K., Sze, S.K., Li, H.Y., Bhushan, S., Wunder, T., and Mueller-Cajar, O. (2023). A linker protein from a red-type pyrenoid phase separates with Rubisco via oligomerizing sticker motifs. Proc Natl Acad Sci U S A. 120: e2304833120.

62. Ohad, I., Siekevitz, P., and Palade, G.E. (1967). Biogenesis of chloroplast membranes. II. Plastid differentiation during greening of a dark-grown algal mutant (Chlamydomonas reinhardi). J Cell Biol. 35: 553–584.

63. Oltrogge, L.M., Chaijarasphong, T., Chen, A.W., Bolin, E.R., Marqusee, S., and Savage, D.F. (2020). Multivalent interactions between CsoS2 and Rubisco mediate α-carboxysome formation. Nature Structural & Molecular Biology. 27: 281–287.

64. Park, J., Bae, S., and Kim, J.-S. (2015). Cas-Designer: a web-based tool for choice of CRISPR-Cas9 target sites. Bioinformatics. 31: 4014–4016.

65. Pettersen, E.F., Goddard, T.D., Huang, C.C., Couch, G.S., Greenblatt, D.M., Meng, E.C., and Ferrin, T.E. (2004). UCSF Chimera--a visualization system for exploratory research and analysis. J Comput Chem. 25: 1605–1612.

66. Punjani, A., Rubinstein, J.L., Fleet, D.J., and Brubaker, M.A. (2017). cryoSPARC: algorithms for rapid unsupervised cryo-EM structure determination. Nat Methods. 14: 290–296.

67. Pyszniak, A.M., and Gibbs, S.P. (1992). Immunocytochemical localization of photosystem I and the fucoxanthin-chlorophyll *a*/*c* light-harvesting complex in the diatom *Phaeodactylum tricornutum*. Protoplasma. 166: 208–217.

68. Ramazanov, Z., Rawat, M., Henk, M.C., Mason, C.B., Matthews, S.W., and Moroney, J.V. (1994). The induction of the CO_2_-concentrating mechanism is correlated with the formation of the starch sheath around the pyrenoid of *Chlamydomonas reinhardtii*. Planta. 195: 210–216.

69. Raven, J.A. (1997). CO_2_-concentrating mechanisms: a direct role for thylakoid lumen acidification? Plant, Cell & Environment. 20: 147–154.

70. Russo, C.J., Dickerson, J.L., and Naydenova, K. (2022). Cryomicroscopy in situ: what is the smallest molecule that can be directly identified without labels in a cell? Faraday Discuss. 240: 277–302.

71. Sander, J.D., and Joung, J.K. (2014). CRISPR-Cas systems for editing, regulating and targeting genomes. Nature Biotechnology. 32: 347–355.

72. Sander, J.D., Maeder, M.L., Reyon, D., Voytas, D.F., Joung, J.K., and Dobbs, D. (2010). ZiFiT (Zinc Finger Targeter): an updated zinc finger engineering tool. Nucleic Acids Research. 38: W462–W468.

73. Schaffer, M., Mahamid, J., Engel, B.D., Laugks, T., Baumeister, W., and Plitzko, J.M. (2017). Optimized cryo-focused ion beam sample preparation aimed at in situ structural studies of membrane proteins. J Struct Biol. 197: 73–82.

74. Schindelin, J., Arganda-Carreras, I., Frise, E., Kaynig, V., Longair, M., Pietzsch, T., Preibisch, S., Rueden, C., Saalfeld, S., Schmid, B., Tinevez, J.Y., White, D.J., Hartenstein, V., Eliceiri, K., Tomancak, P., and Cardona, A. (2012). Fiji: an open-source platform for biological-image analysis. Nat Methods. 9: 676–682.

75. Shimakawa, G., Okuyama, A., Harada, H., Nakagaito, S., Toyoshima, Y., Nagata, K., and Matsuda, Y. (2023). Pyrenoid-core CO_2_-evolving machinery is essential for diatom photosynthesis in elevated CO_2_. Plant Physiology.

76. Shively, J.M., Ball, F., Brown, D.H., and Saunders, R.E. (1973). Functional organelles in prokaryotes: polyhedral inclusions (carboxysomes) of Thiobacillus neapolitanus. Science. 182: 584–586.

77. Sievers, F., Wilm, A., Dineen, D., Gibson, T.J., Karplus, K., Li, W., Lopez, R., McWilliam, H., Remmert, M., Söding, J., Thompson, J.D., and Higgins, D.G. (2011). Fast, scalable generation of high-quality protein multiple sequence alignments using Clustal Omega. Molecular Systems Biology. 7: 539.

78. Smetacek, V. (1999). Diatoms and the ocean carbon cycle. Protist. 150: 25–32.

79. Suchanek, M., Radzikowska, A., and Thiele, C. (2005). Photo-leucine and photo-methionine allow identification of protein-protein interactions in living cells. Nat Meth. 2: 261–268.

80. Tachibana, M., Allen, A.E., Kikutani, S., Endo, Y., Bowler, C., and Matsuda, Y. (2011). Localization of putative carbonic anhydrases in two marine diatoms, *Phaeodactylum tricornutum* and *Thalassiosira pseudonana*. Photosynthesis Research. 109: 205–221.

81. Trifinopoulos, J., Nguyen, L.-T., von Haeseler, A., and Minh, B.Q. (2016). W-IQ-TREE: a fast online phylogenetic tool for maximum likelihood analysis. Nucleic Acids Research. 44: W232–W235.

82. Tsuji, Y., Nakajima, K., and Matsuda, Y. (2017). Molecular aspects of the biophysical CO_2_-concentrating mechanism and its regulation in marine diatoms. Journal of Experimental Botany. 68: 3763–3772.

83. Turk, M., and Baumeister, W. (2020). The promise and the challenges of cryo-electron tomography. FEBS Letters. 594: 3243–3261.

84. Uwizeye, C., Decelle, J., Jouneau, P.-H., Flori, S., Gallet, B., Keck, J.-B., Bo, D.D., Moriscot, C., Seydoux, C., Chevalier, F., Schieber, N.L., Templin, R., Allorent, G., Courtois, F., Curien, G., Schwab, Y., Schoehn, G., Zeeman, S.C., Falconet, D., and Finazzi, G. (2021). Morphological bases of phytoplankton energy management and physiological responses unveiled by 3D subcellular imaging. Nature Communications. 12: 1049.

85. Vernette, C., Lecubin, J., Sánchez, P., Coordinators, T.O., Sunagawa, S., Delmont, T.O., Acinas, S.G., Pelletier, E., Hingamp, P., and Lescot, M. (2022). The Ocean Gene Atlas v2.0: online exploration of the biogeography and phylogeny of plankton genes. Nucleic Acids Research. 50: W516–W526.

86. Villar, E., Vannier, T., Vernette, C., Lescot, M., Cuenca, M., Alexandre, A., Bachelerie, P., Rosnet, T., Pelletier, E., Sunagawa, S., and Hingamp, P. (2018). The Ocean Gene Atlas: exploring the biogeography of plankton genes online. Nucleic Acids Research. 46: W289–W295.

87. Wan, W., and Briggs, J.A. (2016). Cryo-Electron Tomography and Subtomogram Averaging. Methods Enzymol. 579: 329–367.

88. Wang, H., Yan, X., Aigner, H., Bracher, A., Nguyen, N.D., Hee, W.Y., Long, B.M., Price, G.D., Hartl, F.U., and Hayer-Hartl, M. (2019). Rubisco condensate formation by CcmM in β-carboxysome biogenesis. Nature. 566: 131–135.

89. Wunder, T., Cheng, S.L.H., Lai, S.-K., Li, H.-Y., and Mueller-Cajar, O. (2018). The phase separation underlying the pyrenoid-based microalgal Rubisco supercharger. Nature Communications. 9: 5076.

90. Yamashita, K., Palmer, C.M., Burnley, T., and Murshudov, G.N. (2021). Cryo-EM single-particle structure refinement and map calculation using Servalcat. Acta Crystallogr D Struct Biol. 77: 1282–1291.

91. Zang, K., Wang, H., Hartl, F.U., and Hayer-Hartl, M. (2021). Scaffolding protein CcmM directs multiprotein phase separation in β-carboxysome biogenesis. Nat Struct Mol Biol. 28: 909–922.

92. Zhang, K. (2016). Gctf: Real-time CTF determination and correction. J Struct Biol. 193: 1–12.

93. Zheng, S.Q., Palovcak, E., Armache, J.-P., Verba, K.A., Cheng, Y., and Agard, D.A. (2017). MotionCor2: anisotropic correction of beam-induced motion for improved cryo-electron microscopy. Nat Meth. 14: 331–332.

94. Zivanov, J., Nakane, T., Forsberg, B.O., Kimanius, D., Hagen, W.J., Lindahl, E., and Scheres, S.H. (2018). New tools for automated high-resolution cryo-EM structure determination in RELION-3. Elife. 7.

